# Comparative transcriptomics of lateral hypothalamic cell types reveals conserved growth hormone-tachykinin dynamics in feeding

**DOI:** 10.1101/2025.07.28.667087

**Authors:** Vindhya Chaganty, Ruey-Kuang Cheng, Kimberle Shen, Na Zhang, Gaynah J Doblado, Sarah Ong, Sandra Tan, Valarie Yu Yan Tham, Jung-Hwa Choi, Marnie E Halpern, Wei Leong Chew, Anand Kumar Andiappan, Sarah X Luo, Caroline L Wee

## Abstract

The lateral hypothalamus (LH) is a highly heterogeneous brain region regulating hunger and motivated behaviors. In zebrafish, the LH shows distinct neural activity across hunger, feeding, and satiety states. However, the functional and evolutionary conservation of relevant neural circuits remain unclear. Using integrative transcriptomics of zebrafish and mouse LH, we identify conserved cellular clusters with shared molecular markers, particularly within GABAergic neurons. We highlight a conserved GABAergic population expressing tachykinin and growth hormone receptors, which is responsive to food cues and modulated by hunger and feeding states. This cluster may mediate acute appetite-enhancing effects of growth hormone. In both species, feeding elevates growth hormone receptor and tachykinin expression and activates these neurons, while human growth hormone increases their activity and food intake in zebrafish. These findings suggest a conserved neural mechanism by which metabolic hormones influence feeding behavior. Our comparative LH atlas highlights the evolutionary biology of appetite regulation and the integration of hormonal and neural signals driving energy homeostasis.

## Introduction

Central neural circuits, especially within the hypothalamus, are crucial for maintaining the balance between energy intake and expenditure, by integrating environmental and internal metabolic cues (*1–3*). The lateral hypothalamus (LH) / lateral hypothalamic area (LHA) and ventromedial hypothalamus (VMH) have been studied extensively in the context of appetite regulation (*4–7*). Both lesioning and stimulation experiments in mammals have established the LHA and VMH as the “hunger” and “satiety” centers respectively (*8–13*). As in mammals, larval zebrafish exhibit complex feeding behaviors and contain brain regions and neuromodulatory circuits implicated in appetite control. Recently, the inferior lobe of the ventral hypothalamus was proposed as the zebrafish counterpart of the mammalian LH (*14*, *15*), where it was shown to be activated both by the sensory perception and consumption of food. LH neurons show anti-correlated activity with the ventromedial caudal hypothalamus (cH) drawing parallels for what has been observed in the mammalian VMH (*16*, *17*). Further, circuit manipulation studies demonstrated functional connectivity between the two regions, suggesting inhibitory crosstalk between the LH and cH (*14*).

The results suggest that these zebrafish hypothalamic regions have energy and nutrient sensing capabilities, but several questions remain. First, the overall cellular and molecular identities and functions of their cell types are unknown; this is especially so for the highly heterogeneous LH (*18*). For example, hypocretin and MCH-expressing neurons characteristic of mammalian LH are not found in the zebrafish LH but rather located in close proximity (*14*). One study has even proposed that the zebrafish inferior lobe (LH) has a mesencephalic rather than diencephalic origin, raising questions regarding the evolutionary homology of this region (*19*).

Further, while GABAergic and glutamatergic populations have been shown to enhance or suppress feeding respectively in rodents (*20*) and show remodeling during obesity (*21*), more specific cellular subtypes in the mouse LH have been described but not functionally characterized (*18*, *22*), and their degree of conservation in non-mammalian vertebrate species is unknown. The molecular comparison of LH cell types between teleost fish and mammals and the identification of conserved functional populations would allow a more comprehensive understanding of the diversity of functions of the LH in the light of evolution.

Here, we generated a zebrafish LH scRNAseq dataset based on an established transgenic marker, and then using the Act-seq method to preserve neuronal activity signatures, we pinpointed specific cellular subtypes in the zebrafish LH that were differentially activated during voracious feeding. Then, using an integrative omics approach, we mapped the zebrafish dataset to a published mouse dataset (*18*) to identify both overlapping and distinct cellular clusters. We identified several conserved neuropeptides and receptors, including a growth hormone-tachykinin population that was not previously identified from mouse scRNAseq data clustering alone. Notably, the expression of growth hormone (GH) in the pituitary and its receptor in an LH tachykinin cell type are both dynamically regulated by hunger and satiety. Further, GH administration was able to acutely potentiate feeding in zebrafish. Our findings suggest that beyond its longer-term effects on growth, appetite and metabolism, GH has an acute and direct role in appetite stimulation through activation of neuronal pathways, potentiated in part through the newly identified LH subpopulation of GH receptor-expressing tachykinin neurons. We propose that this conserved hormonally regulated circuit is used to integrate an animal’s appetite with its growth and metabolic needs, so that energy supply adequately meets demand.

## RESULTS

### Act-seq identifies distinct cell types in the LH that are responsive to feeding state

Several studies in mouse models have shown that the LH is highly heterogeneous (*18*, *22–24*) yet much of this heterogeneity and its conservation is not functionally characterized. Hence, we sought to identify specific sub-populations in the zebrafish LH which are conserved in mammals and show hunger and feeding state-dependent changes in activity. To this end, we adopted the Act-seq method (*25*) to preserve feeding stimulus-based activity patterns in neurons by reducing dissociation-induced transcription. Larval zebrafish carrying the *Tg(76A:Gal4FF;UAS:GCaMP6s)* transgene which labels the LH were randomly divided into two groups: “Food-deprived” (for 4 hours) and “Voracious feeding” (food-deprived for 4 hours and then supplied with paramecia, which correlates with enhanced hunting and feeding) (*14*, *26*, *27*). Dissociation was carried out in the presence of Actinomycin D to inhibit transcription (see Methods) (**Figure 1A**). GCaMP6s-positive cells were isolated using FACS and combined with the negative population in roughly equivalent numbers to ensure sufficient capture of LH cells. We surveyed a total of 9031 cells (4412 cells from the “Food-deprived” group and 4619 cells from the “Voracious feeding” group). Cells were then filtered for neuronal identity and divided into LH (*Gal4FF*-positive) and non-LH groups based on the expression of *Gal4FF* (**Figure 1A, Figure S1**). The purpose of including LH-negative cells was both to serve as an internal control for the Act-seq protocol and for comparison to the LH-positive population.

**Figure 1.**
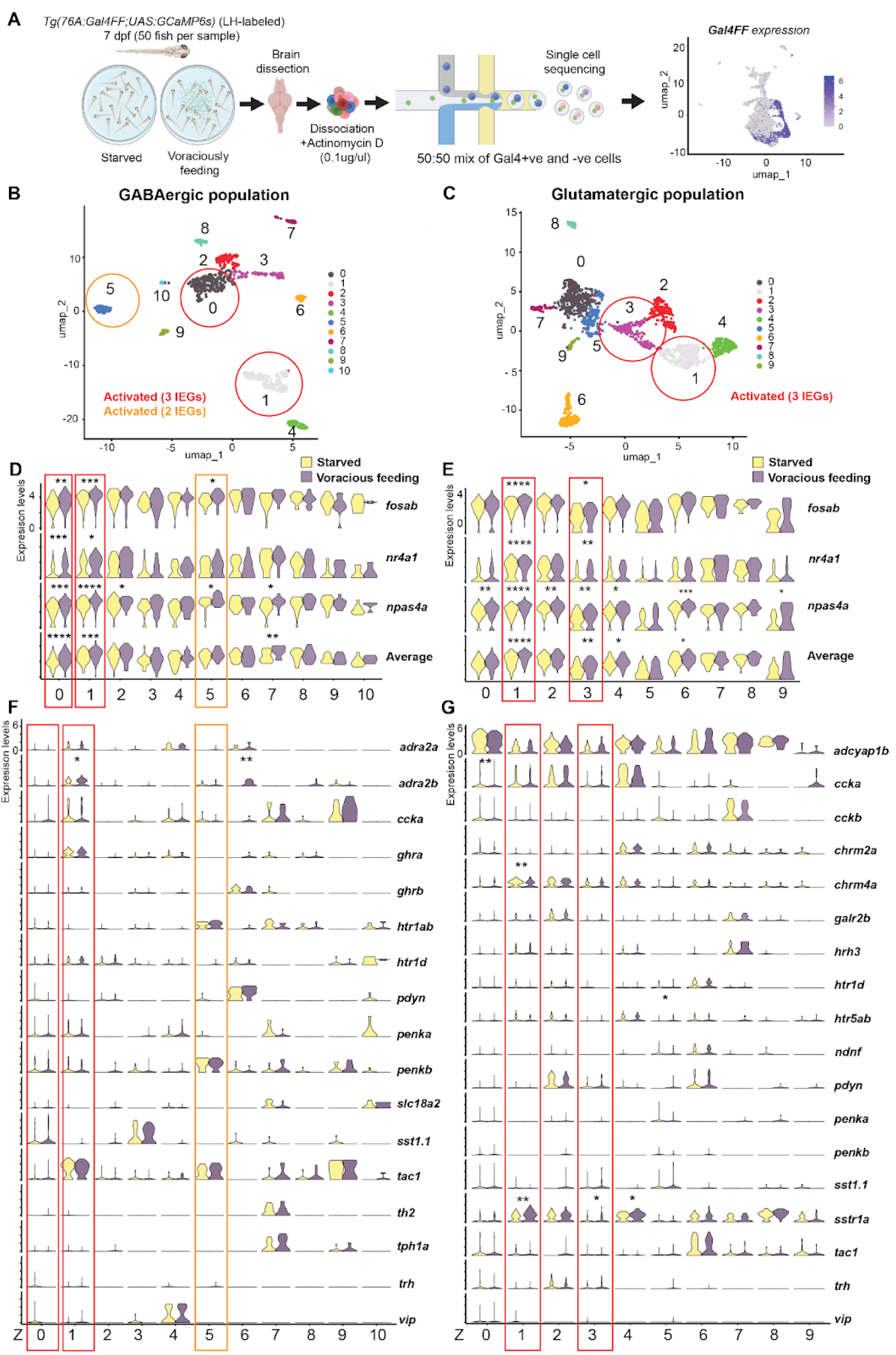
Act-seq identifies a conserved neuronal sub-cluster in the LH activated by voracious feeding. **A)** Schematic of the Act-seq workflow including the identification of LH cells by *Gal4FF* expression. **B)** UMAP of LH cells sub-clustered based on their GABA and **C)** glutamatergic identities. Activated clusters are circled in red (all three Immediate Early Genes (IEGs) show increased expression in voracious feeding relative to food-deprived) or orange (⅔ IEGs show increased expression) **D)** Individual and average IEG expression of each GABAergic or **E)** Glutamatergic cluster in food-deprived (yellow) or voracious feeding (purple) conditions. **F)** Expression of selected neuropeptides and receptors (cluster marker genes or appetite-related genes) in food-deprived (yellow) or voracious feeding (purple) conditions in GABAergic or **G)** glutamatergic populations. Activated clusters are indicated by red (3 IEGs) or orange (2 IEGs) boxes. Significant increases in gene expression across conditions are also indicated with asterisks. *p<0.05, **p<0.01. ***p<0.001

We further subdivided the LH population based on their GABAergic and glutamatergic identities as these populations have been shown to differentially regulate feeding in mammals (*20*, *28*). Sub-clustering GABA (*slc32a1*-positive) and glutamatergic (*slc17a6a*-positive) cell types revealed 11 GABAergic (897 cells) and 10 glutamatergic (1817 cells) clusters (**Figure 1B-C**, **Table 1–2**). We further quantified the activity of each cluster using the relative expression of three Immediate Early Genes (IEGs), *fosab* (commonly known as *c-fos*), *nr4a1*, and *npas4a* (*29, 30*) (**Figure 1D-E**).

**Table 1:** Zebrafish GABAergic clusters and their markers.

| Cluster (asterisks indicated activated clusters) | Markers reported in Mickelsen et al., (2019) (18) dataset (top 60) | Neuromodulator and receptor markers (top 60) |
| --- | --- | --- |
| **Z0<sup>GABA\*\*\*</sup>** | *CABP7, cxxc5a, fstl5, nanos1, NPAS3, nxph1, stmn3, zfhx4* |  |
| **Z1<sup>GABA\*\*\*</sup>** | *adra2a, ghra, gsx1, ngef, prox1a, tac1* | *adra2a, adra2b, cart2, chrm4a, ghra, oprd1b, tac1* |
| **Z2<sup>GABA</sup>** | *cbln2b, fam43b, foxp2, lmo1, lmo4b, luzp2, meis2a, mllt3, nptx1l* | *chrm2a* |
| **Z3<sup>GABA</sup>** | *arxa, KCNIP4, man1a1, necab2, nxph1, six3a, sst1.1, tac3a* | *adra2da, sst1.1, tac3a* |
| **Z4<sup>GABA</sup>** | *adra2a, bhlhe22, cadps2, dlx1a, fign, gsx1, lmo3, nkx2.1, pou6f2, znf536* | *adra2a, npr3, vip* |
| **Z5<sup>GABA\*\*</sup>** | *arxa, cntn5, ctbp2a, kcnab1a, lhx6, nkx2.2a, sp9, sphkap* | *cart2, htr1ab, penkb* |
| **Z6<sup>GABA</sup>** | *hmx2, pdyn, sox5* | *galr1b, galr2b, ghrb, pdyn, ptger4c,sstr5, vipr1b* |
| **Z7<sup>GABA</sup>** | *ARHGAP6, ddc, lef1, slc18a2, th2, zbtb20* | *ddc, scg3, th2, tph1a, slc18a2* |
| **Z8<sup>GABA</sup>** | *abhd16a, cdh11, nkx2.1, nptx1l, sp9* | *chrm2a, drd3, trhr2* |
| **Z9<sup>GABA</sup>** | *nos1, pou3f2b* | *adcyap1b, ccka* |
| **Z10<sup>GABA</sup>** | *cacng5a, ddc, gfra1a, ntsr1, slc18a2* | *ddc, htr1d, ntsr1, tph2, slc18a2* |

**Table 2:** Zebrafish Glutamatergic clusters and their markers.

| Cluster (asterisks indicated activated clusters) | Markers reported in Mickelsen et al., (2019) (18) dataset (top 60) | Neuromodulator and receptor markers (top 60) |
| --- | --- | --- |
| **Z0<sup>Glut</sup>** | *adcyap1b, CABP7, khdrbs2, nanos1, meis2a, msi2a, necab2, NPAS3, pbx3b* | *adcyap1b* |
| **Z1<sup>Glut\*\*\*</sup>** | *cpne2, gpr22a, junba, pip5k1ba* |  |
| **Z2<sup>Glut</sup>** | *cbln4, ccka, lmo3, necab1, nos1, pdyn, pou3f2b, SORCS3* | *ccka, chrm4a, galr2b, pdyn* |
| **Z3<sup>Glut\*\*\*</sup>** | *lmo3, prdm8b* |  |
| **Z4<sup>Glut</sup>** | *ccka, doc2a, epha4a, gpr22a, KHDRBS3, hopx, SYT2* | *ccka, chrm2a, htr5ab* |
| **Z5<sup>Glut</sup>** | *meis2a, nanos1, nrxn3a, pbx3b* | *gad2* |
| **Z6<sup>Glut</sup>** | *camkva, cbln1, fezf1, gabra5, hlfb, kctd12.1, sox1a, sox2, sox5, tac1* | *chrm2a, htr1d, ndnf, tac1* |
| **Z7<sup>Glut</sup>** | *bhlhe22, gabra5, gem, mpped2, neurod2, prdm8b, sema3e, tbr1b, vim* | *cckb, hrh3, prlhr2a* |
| **Z8<sup>Glut</sup>** | *adka, cadps2, caln1, gfra1a, khdrbs2, necab2, pou3f2b, rftn1a, sez6a, thsd7ba, tshz1* | *sstr1a* |
| **Z9<sup>Glut</sup>** | *b3gat2, lmo2, neurod1, neurod2, pax6a, zic1, zic4* | *ndnf* |

As expected, many LH clusters expressed neuropeptide-related genes such as *tac1, sst, th, slc18a2, cck, trh*, *penk, pdyn,* and *vip*, as well as neuroreceptors for growth hormone, monoamines (serotonin, dopamine, norepinephrine), and acetylcholine (**Figure 1F-G**, **Table 1-2**), many of which are also present in the mouse LH (*18*). In general, the GABAergic population had a larger number of neuromodulators and receptors within their top marker genes (transcripts that are highly expressed in a specific cell type and distinguish them from other cell types) (**Table 1-2)**. These neuropeptides and receptors were also present in adult zebrafish LH, as demonstrated by bulk transcriptomics analyses performed in parallel (**Figure S2**).

Of the activated GABAergic clusters, two zebrafish GABAergic clusters 0 and 1 (which we will refer to as Z0^GABA^ and Z1^GABA^) showed increased expression of all three IEGs, and one (Z5^GABA^) showed increased expression of two IEGs in voracious feeding condition when compared to the starved condition (**Figure 1D**). Z0^GABA^ expresses marker genes such as *CABP7, cxxc5a, fstl5, nanos1, NPAS3, nxph1, stmn3, zfhx4,* which were also mouse LH marker genes reported in (*18*), with no neuromodulators or receptors appearing within the top 60 marker genes (**Table 1**). Z1^GABA^ on the other hand expressed a variety of neuromodulator and receptor markers such as *tac1* (tachykinin 1)*, ghra* (growth hormone receptor a)*, cart2, adra2a, adra2b, chrm4a,* and *oprd1b*, with *ghra* and *chrm4a* also appearing as the top 5 marker genes. Z5^GABA^ expressed *penkb, htr1ab,* and *cart2* (**Figure 1F**, **Table 1**).

Two glutamatergic clusters showed increased expression of all three IEGs (**Figure 1E**). Zebrafish glutamatergic Cluster 1 (which we will refer to as Z1^Glut^) expressed markers such as *cpne2, gpr22a, junba, pip5k1ba* (**Table 2**) and Z3^Glut^ expressed *lmo3, prdm8b*, all of which were previously reported in mouse LH clusters (*18*). Neither cluster had neuromodulators or receptors within their top 60 marker genes (**Table 2**), although their neurons do express neuromodulators and receptors such as *adcyap1b*, *ccka*, *chrm4a*, and *sstr1a* which are expressed in multiple glutamatergic clusters (**Figure 1G**).

We mapped the spatial expression patterns of some key neuropeptides in the zebrafish LH using Hybridisation chain reaction (HCR) (**Figure 2A**) and also examined neuronal activity through co-expression of *fosab* expression (*29*). We performed this analysis across three feeding states, “food-deprived” (for 4 hours), “voracious feeding” (food-deprived for four hours and then fed paramecia) and “fed (satiated)”, in which larvae were continuously fed with excess paramecia. Our analyses revealed cell-type specific and feeding state-dependent dynamics (**Figure 2B-C**).

**Figure 2.**
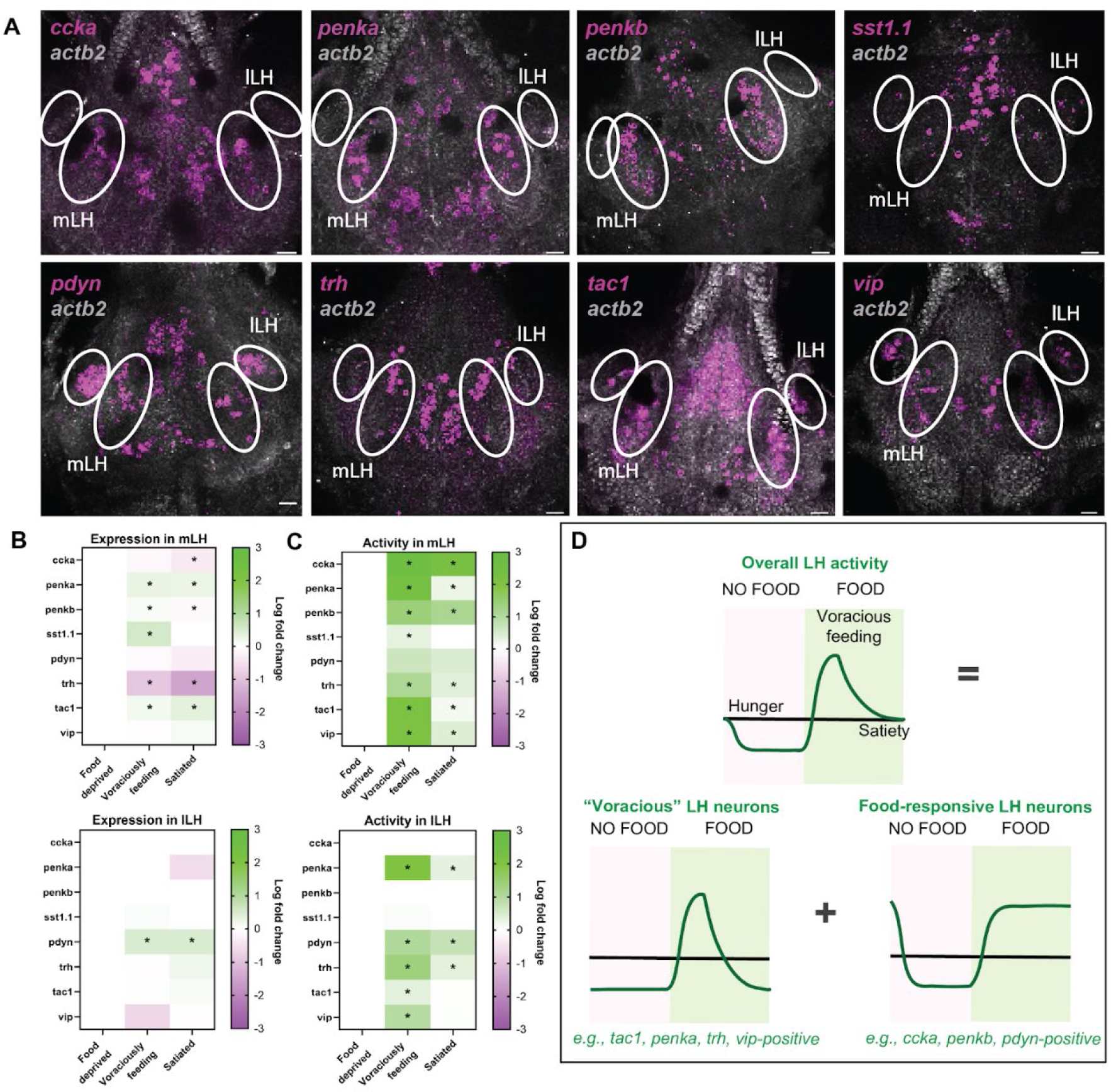
Conserved receptor and neuromodulator expression in zebrafish larval hypothalamic regions. **A)** Representative images showing expression of important neuromodulators (*ccka, penka, penkb, sst1.1, pdyn, trh, tac1, vip*) in 7 dpf larval brains using HCR. **B)** Heatmaps showing dynamic expression of neuromodulators in medial lateral hypothalamus (mLH, top) and lateral lateral hypothalamus (lLH, bottom) respectively under voraciously feeding and satiated conditions with respect to food deprived conditions. The mLH and lLH were analysed separately as they may have different functional roles in sensory versus consummatory feeding behaviour (Wee et al., (2019). **C)** Heatmaps showing dynamic activity reflected by *fosab* expression of neuromodulator expressing neurons under different feeding states with respect to food deprived conditions (mLH, top, lLH, bottom). Expression and activity values were averaged from individual neurons from 8-10 fish per condition per gene. Genes with significant differential expression or activity as compared to the food deprived state are labeled with an asterisk (FDR < 0.05, Statistical analysis performed using unpaired t-test with Welch correction). **D)** Schematic showing how activity patterns of different neuropeptidergic populations may reflect the composite LH activity patterns reported in Wee et al., (2019).

Neurons expressing *penka, sst1.1, tac1* and *vip* were most active during voracious feeding, whereas their activity was reduced in fed (satiated) fish. We hypothesize that these neurons could be sensitized by the cH (*14*) or other hunger hormones during food deprivation and they play a role in the subsequent upregulation of feeding (i.e., voracious feeding). In contrast, neurons expressing *ccka, penkb, pdyn* and *trh* are active upon food presentation and remain active even in fed (satiated) larvae. These neuronal subtypes may encode the presence of food and could also be satiety-sensing neurons that promote satiation. Together, the composite activity patterns of these two broad classes of neurons can explain the overall LH activity patterns across feeding states (**Figure 2D**).

Of the non-LH clusters, glutamatergic and GABAergic neurons (all 3 IEGs), as well as cranial ganglia (2 IEGs) showed significantly higher activity under voracious feeding, with retina cells showing higher *fosab* expression **(Figure S1A and B).** We further sub-clustered the GABAergic **(Figure S1C and D)** and glutamatergic **(Figure S1E and F)** non-LH neurons and identified activated clusters including an *orthopedia a* (*otpa*)*-* expressing GABAergic cluster and glutamatergic cerebellar granule cells. These results are consistent with the general enhancement of visuomotor activity associated with voracious feeding and confirm the robustness of the Act-seq analysis.

### Integration of zebrafish and mouse LH transcriptomes uncovers both conserved and distinct cell types

A systematic comparison of zebrafish and mouse LH has not previously been performed. We hence sought to compare our zebrafish clusters to a mouse dataset (*18*). Independent UMAP (Uniform Manifold Approximation and Projection for Dimension Reduction) analyses of the zebrafish and mouse LH datasets revealed multiple GABAergic and glutamatergic clusters that are similar across the two species, based on the expression of conserved marker genes (**Figure 3, Figure S3**).

To further examine cross-species homology, we applied an integrative omics approach for cross-species mapping of GABA and glutamatergic populations into a shared embedding space (**Figure 4**), applying iterative clustering using the Seurat canonical correlation analysis (CCA) workflow (*31*). The integrated GABAergic neurons (**Figure 4A-C, G, Table 3**) clustered into 12 cell types, with strong representation of both zebrafish and mouse neurons within each cluster, except for Integrated GABAergic Clusters 10 and 11 (Int10^GABA^and Int11^GABA^) which comprised mainly zebrafish neurons and Int9^GABA^ which comprised mainly mouse neurons. In contrast, 10 of the 16 integrated glutamatergic clusters comprised either predominantly zebrafish neurons (Int3^Glut^, Int6^Glut,^ Int7^Glut^, Int9^Glut^, Int13^Glut^) or mouse neurons (Int8^Glut^, Int12^Glut^, Int14^Glut^, Int15^Glut^). Only Int0^Glut^, Int1^Glut^, Int2^Glut^, Int4^Glut^, Int5^Glut^, Int10^Glut^, and Int11^Glut^ contained both zebrafish and mouse neurons (**Figure 4D-F, H, Table 4**). These results suggest stronger conservation of GABAergic rather than glutamatergic LH cell types.

**Table 3:**
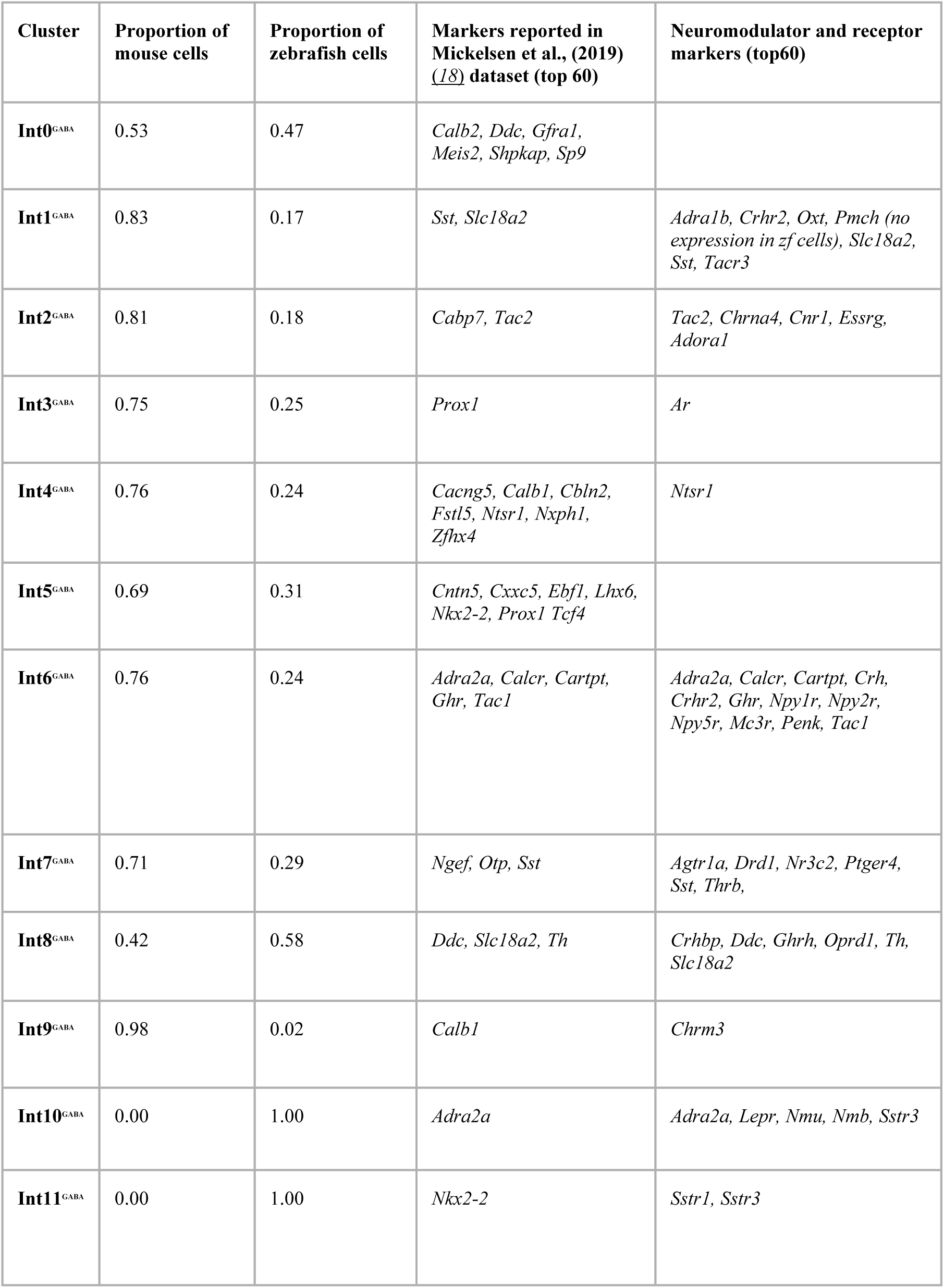
Integrated GABAergic clusters and their markers.

| Cluster | Proportion of mouse cells | Proportion of zebrafish cells | Markers reported in Mickelsen et al., (2019) (18) dataset (top 60) | Neuromodulator and receptor markers (top60) |
| --- | --- | --- | --- | --- |
| **Int0<sup>GABA</sup>** | 0.53 | 0.47 | *Calb2, Ddc, Gfra1, Meis2, Shpkap, Sp9* |  |
| **Int1<sup>GABA</sup>** | 0.83 | 0.17 | *Sst, Slc18a2* | *Adra1b, Crhr2, Oxt, Pmch (no expression in zf cells), Slc18a2, Sst, Tacr3* |
| **Int2<sup>GABA</sup>** | 0.81 | 0.18 | *Cabp7, Tac2* | *Tac2, Chrna4, Cnr1, Essrg, Adora1* |
| **Int3<sup>GABA</sup>** | 0.75 | 0.25 | *Prox1* | *Ar* |
| **Int4<sup>GABA</sup>** | 0.76 | 0.24 | *Cacng5, Calb1, Cbln2, Fstl5, Ntsr1, Nxph1, Zfhx4* | *Ntsr1* |
| **Int5<sup>GABA</sup>** | 0.69 | 0.31 | *Cntn5, Cxxc5, Ebf1, Lhx6, Nkx2-2, Prox1 Tcf4* |  |
| **Int6<sup>GABA</sup>** | 0.76 | 0.24 | *Adra2a, Calcr, Cartpt, Ghr, Tac1* | *Adra2a, Calcr, Cartpt, Crh, Crhr2, Ghr, Npy1r, Npy2r, Npy5r, Mc3r, Penk, Tac1* |
| **Int7<sup>GABA</sup>** | 0.71 | 0.29 | *Ngef, Otp, Sst* | *Agtr1a, Drd1, Nr3c2, Ptger4, Sst, Thrb,* |
| **Int8<sup>GABA</sup>** | 0.42 | 0.58 | *Ddc, Slc18a2, Th* | *Crhbp, Ddc, Ghrh, Oprd1, Th, Slc18a2* |
| **Int9<sup>GABA</sup>** | 0.98 | 0.02 | *Calb1* | *Chrm3* |
| **Int10<sup>GABA</sup>** | 0.00 | 1.00 | *Adra2a* | *Adra2a, Lepr, Nmu, Nmb, Sstr3* |
| **Int11<sup>GABA</sup>** | 0.00 | 1.00 | *Nkx2-2* | *Sstr1, Sstr3* |

**Table 4:**
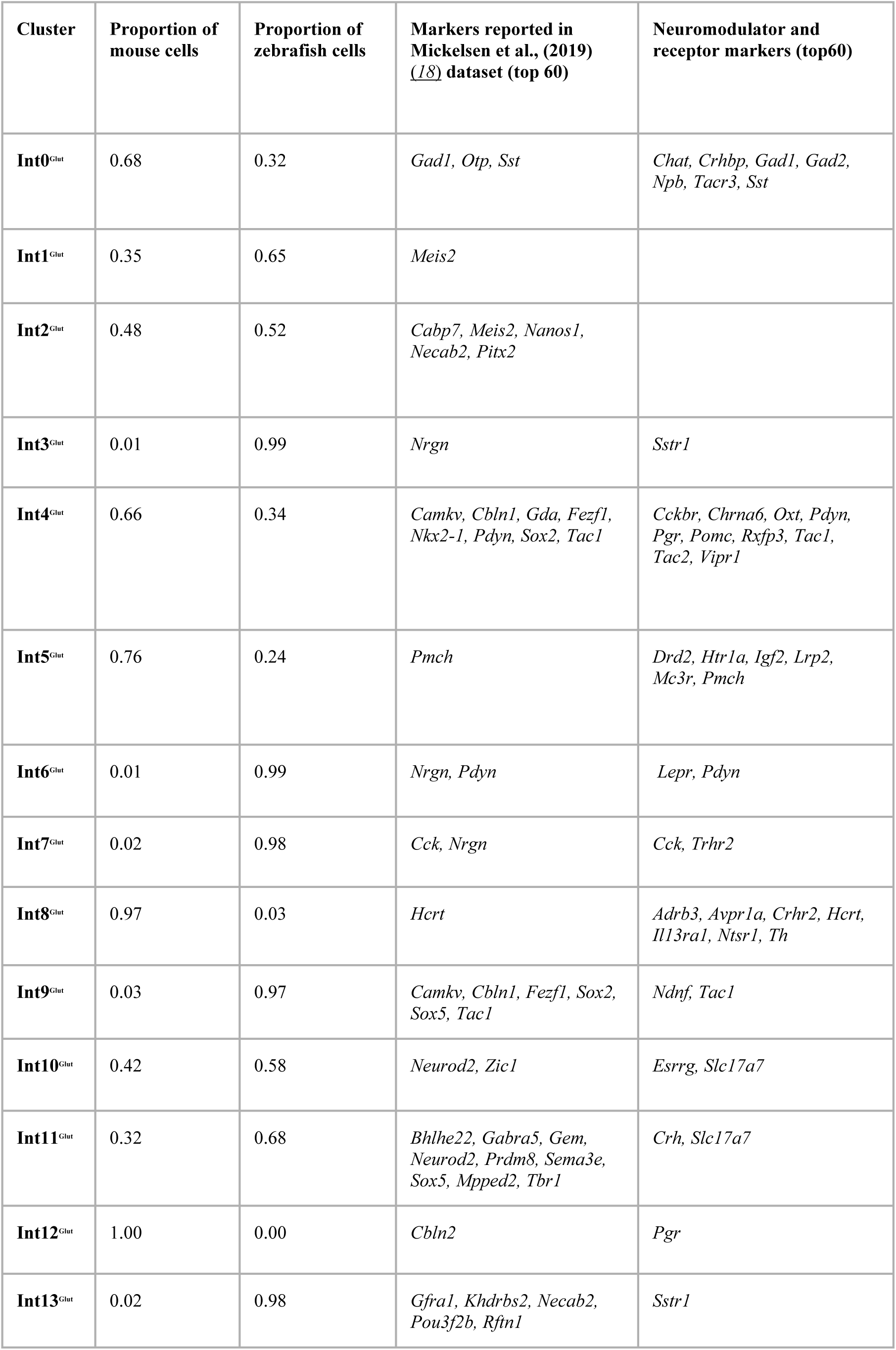
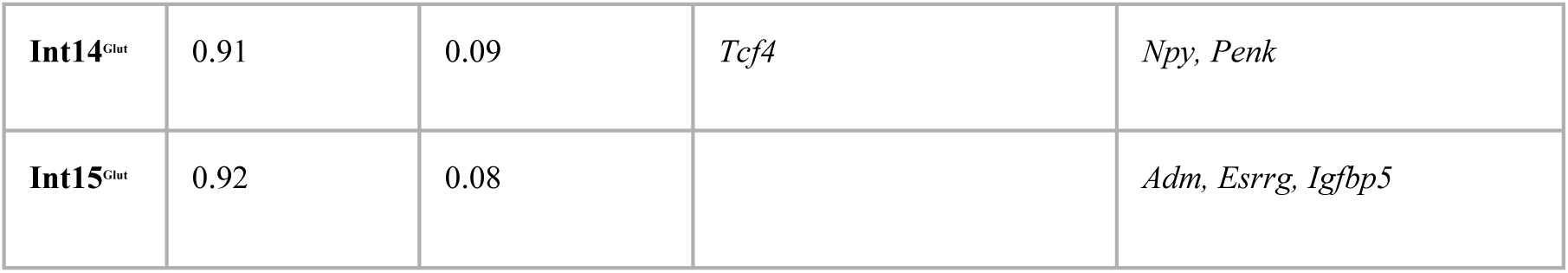
Integrated Glutamatergic clusters and their markers.

It has been shown previously that zebrafish LH does not consist of some landmark neuronal populations for eg., neurons expressing hypocretin/orexin (specified by the *hcrt* gene) and melanin concentrating hormone produced by the *pmch* gene (*14*). As expected, since zebrafish *hcrt* is not expressed in the LH, *Hcrt* was a marker for the mouse-specific Int8^Glut^ and though some *hcrt-*expressing cells were distributed into other integrated clusters, they were still restricted to mouse neurons within these clusters (**Figure 4H**). Similarly, *Pmch* showed up as a cluster marker in Int5^Glut^ (which comprised mainly mouse neurons), and its expression is only restricted to mouse neurons in this cluster. As described in (*18*), which generated the mouse dataset, low levels of *Pmch* expression were found in all glutamatergic and GABAergic neurons, which was attributed to ambient mRNA released from damaged neurons during dissociation. There were also other significant differences between zebrafish and mouse LH clusters, for example many more zebrafish LH clusters expressed *adcyap1b*, which was absent in most mouse clusters (**Figure 1F-G**, **Figure 3B, D, Table 1–2**). Besides *Hcrt* and *Pmch*, the mouse LH also expressed neuromodulatory markers that were not found in fish LH, such as *Nts* and *Gal* in GABAergic neurons (**Figure 3B**).

**Figure 3.**
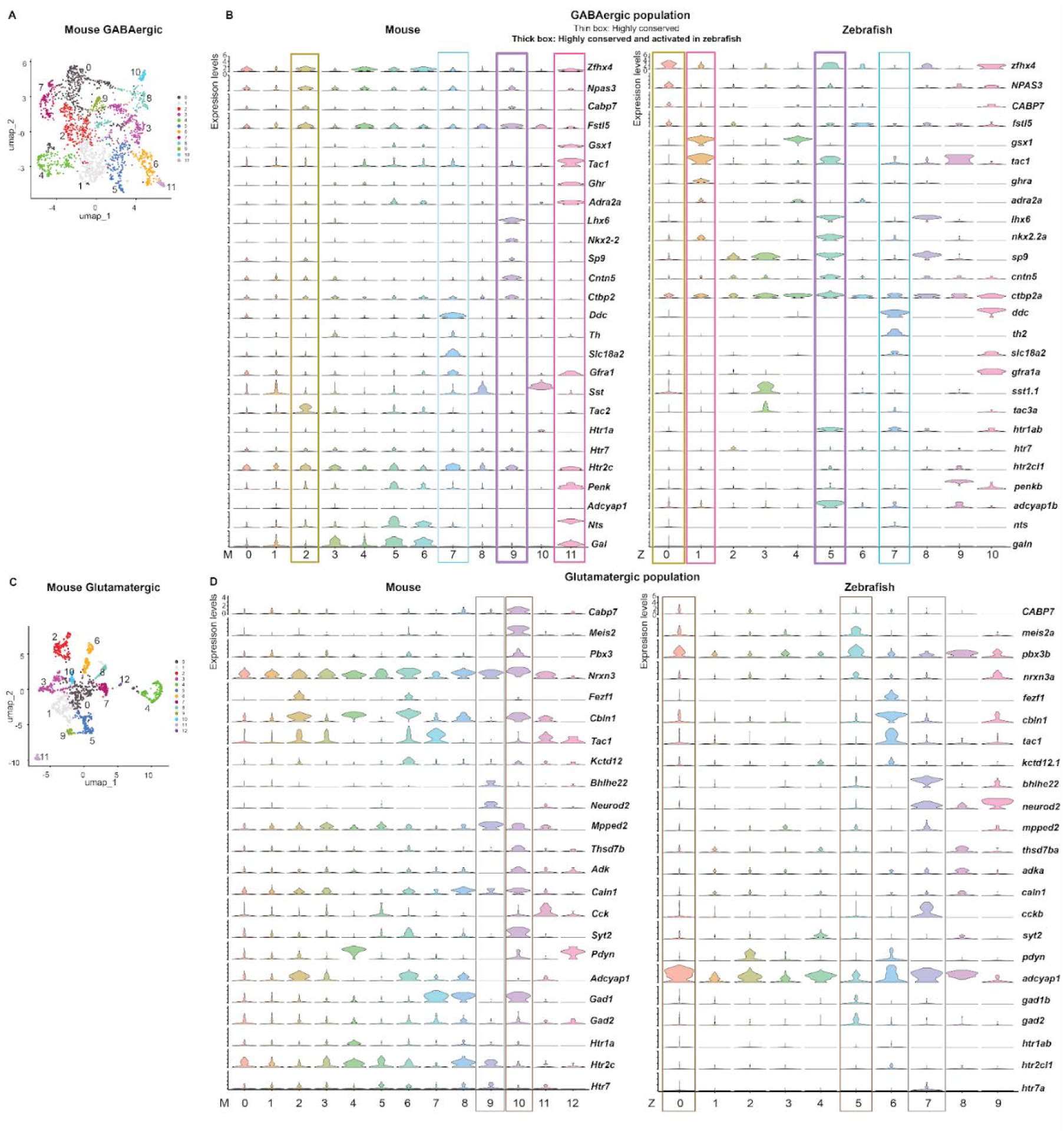
Inter-species comparison of mouse and zebrafish LH transcriptomes. a) UMAP of mouse LH sub-clustered by **A)** GABAergic and **C)** glutamatergic identities **B)** Common markers expressed in mouse and zebrafish GABAergic and **D)** glutamatergic clusters. Clusters with similar expression of marker genes are indicated by the same-coloured boxes. Clusters that are highly conserved in both species are indicated by thin boxes and clusters that are highly conserved and also activated in zebrafish are highlighted with thick boxes. Note that they are all GABAergic. GABAergic: M2^GABA^ and Z0^GABA^ (yellow); M7^GABA^ and Z7^GABA^ (blue); M9^GABA^ and Z5^GABA^ (purple); M11^GABA^ and Z1^GABA^ (pink). Glutamatergic: M9^Glut^ and Z7^Glut^ (grey); M10^Glut^ and both Z0^Glut^ and Z5^Glut^ (brown).

**Figure 4.**
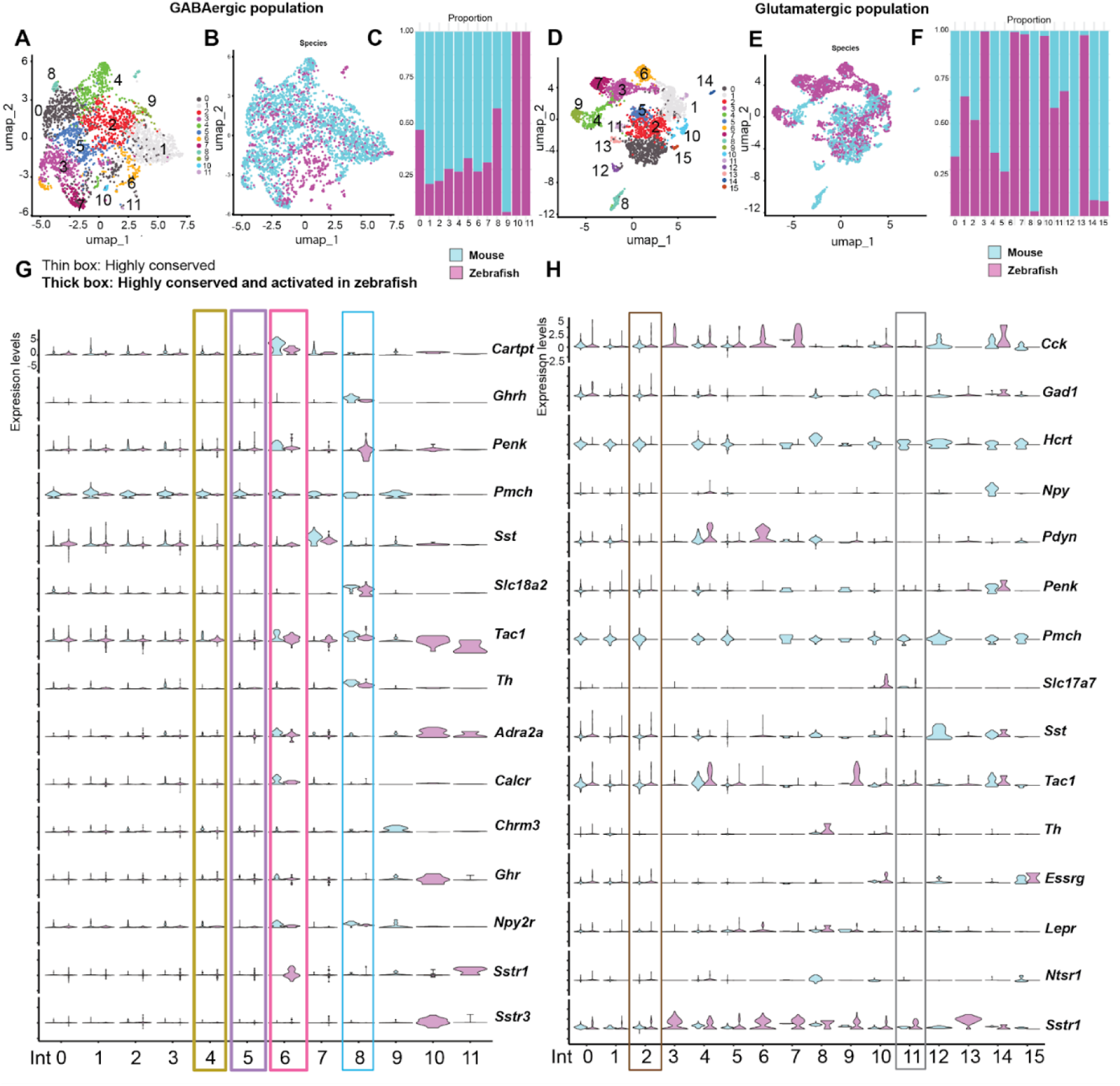
Single-cell transcriptomic integration of mouse and zebrafish LH. Cross-species integration of zebrafish and mouse LH (*18*) using the Seurat CCA Integration tool. UMAP visualisation of the integrated dataset sub-clustered by **A)** GABAergic and **D)** glutamatergic identities. Species specific distribution of mouse (cyan) and zebrafish (magenta) cells in **B)** GABAergic and **E)** glutamatergic populations. Proportions of mouse (cyan) versus zebrafish (magenta) cells in **C)** GABAergic and **F)** glutamatergic populations. Expression of selected neuropeptides and receptors in mouse (cyan) or zebrafish (magenta) cells in the **G)** integrated GABAergic or **H)** integrated glutamatergic populations. Color code of outlined boxes is shared with Figure 3. Clusters that are highly conserved in both species are indicated by thin boxes and clusters that are highly conserved and also activated in zebrafish are highlighted with thick boxes.

We did not observe clear similarities between the activated zebrafish glutamatergic clusters (Z1^Glut^ and Z3^Glut^) with the integrated or mouse clusters. However, Z0^Glut^ and Int2^Glut^ shared marker genes *Cabp7, Meis2, Nanos1,* and *Necab2*, and was most similar to M10^Glut^ which had markers *Cabp7*, *Meis2* and *Pbx3* overlapping with Z0^Glut^ (**Figure 3D, Table 4 and 5**). *gad2*-positive Z5^Glut^ also showed some overlap with M10^Glut^, sharing markers *Meis2, Pbx3, Nrxn3* (**Figure 3D, 4H, Table 4 and 5**). Notably, both Z5^Glut^ and M10^Glut^ express *gad1b/Gad1* and *gad2/Gad2* (**Figure 3D**), suggesting conservation of dual glutamatergic and GABAergic cellular identities, as also previously reported in (*18*). Z7^Glut^, M9^Glut^, and Int11^Glut^ shared marker genes *Bhlhe22*, *Neurod2*, and *Mpped2* (Figure 3D, Table 4 and 5).

On the other hand, some of the integrated clusters, especially GABAergic ones, showed stronger conservation of marker genes in both zebrafish and mouse neurons. In particular, Int8^GABA^ co-expresses *Th*, *Ddc,* and *Slc18a2* (**Figure 4G**, **Table 3 and 5**), which are markers for dopamine synthesis (*Th*, encodes for Tyrosine hydroxylase, *Ddc*, encodes for Dopamine decarboxylase), and transport (*Slc18a2*, encodes Vesicular monoamine transporter 2 or Vmat2). It is also highly similar to Mouse GABAergic Cluster 7 (M7^GABA^) ^and^ Z7^GABA^ (Z10^GABA^ to a smaller extent, which also expresses *ddc, gfra1* and *slc18a2*) (**Figure 1F, 3B, Table 3 and 5**). This cluster was also identified by Mickelsen et al., (2019) (*18*) and we confirmed the co-expression of *th2* and *slc18a2* in the zebrafish LH using immunostaining (**Figure S4B**).

Furthermore, all the GABAergic clusters identified to be activated in zebrafish also had a corresponding cluster in the integrated dataset suggesting a conserved role across the two species. Z5^GABA^, which had activation of 2 out of 3 IEGs, showed closest overlap with Int5^GABA^ (and shares the markers *Lhx6, Nkx2-2, Sp9, Cntn5* and *Ctbp2* with M9^GABA^), with common overlap of *Lhx6, Nkx2-2,* and *Cntn5* in the integrated cluster (**Figure 3B**, **Table 3 and 5**). *penkb* and *htr1ab,* which are markers of Z5^GABA^ are also expressed in this M9^GABA^ cluster (*Penk* and *Htr1a)*, albeit at lower levels (**Figure 3B**, **Table 1**).

Of the two highly activated zebrafish LH clusters (increased expression of all 3 IEGs expression), Z0^GABA^ showed close similarity to Int4^GABA^ (and shares the markers *Zfhx4*, *Npas3*, *Cabp7*, and *Fstl5* with M2^GABA^) (**Figure 3B**, **Table 3 and 5**). Notably, Z1^GABA^, showed high similarity to both Int6^GABA^ and M11^GABA^, sharing a common set of neuropeptide and receptors (*Tac1*, *Ghr, Adra2a*) as well as other markers (*Gsx1*) (**Figure 3B, 4G, Table 3 and 5)**. We will refer to this Z1^GABA^ cluster as the *tac1/ghra* cluster in the next section. Overall, these results highlight that all activated GABAergic clusters discovered in fish also have clear counterparts in mammals. A summary of all conserved clusters described above can be also found in **Table 5**.

**Table 5:**
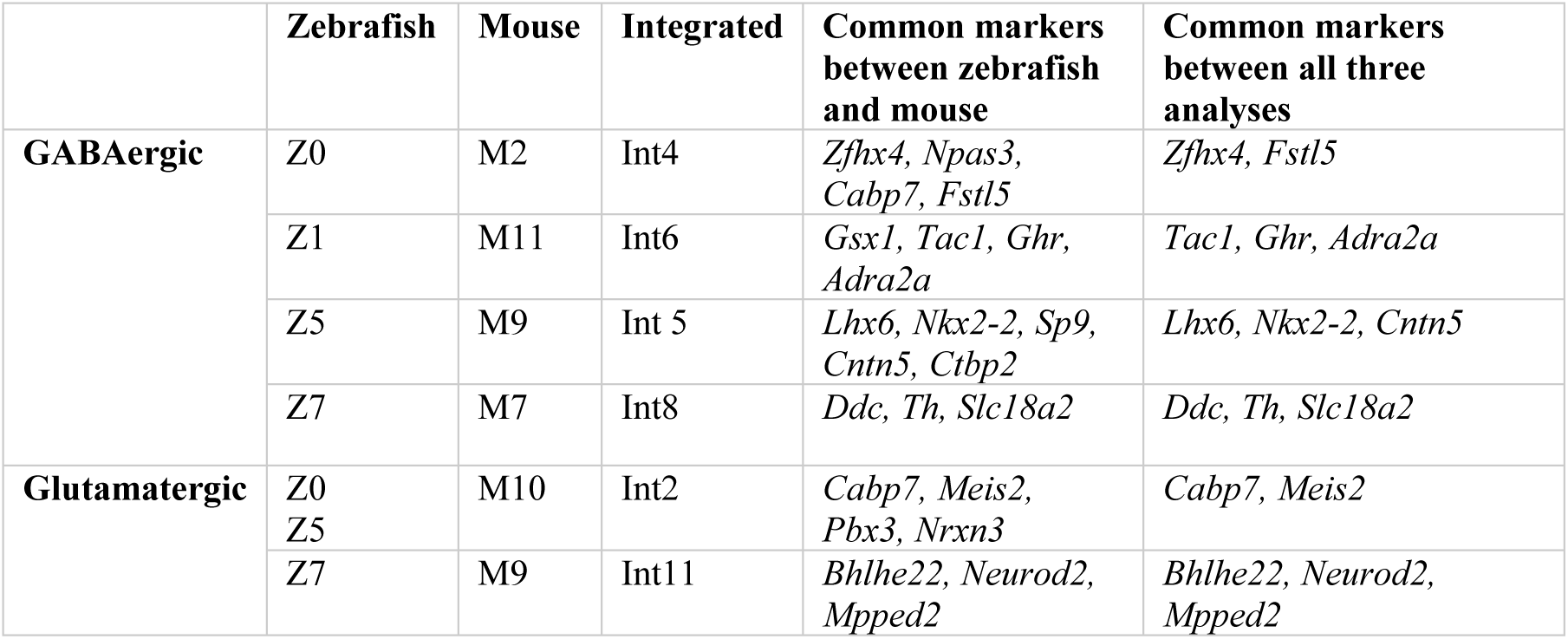
Comparison of conserved clusters and their markers.

### Zebrafish *tac1/ghra* neurons show dynamic activity and expression across feeding states

The Act-seq data and integrative comparative omics strongly highlights the GABAergic *tac1/ghra* population as a highly conserved LH cell type which showed significantly higher expression of all three IEGs during voracious feeding. Interestingly, the density of growth hormone receptor transcripts (*ghra*) was highest in the LH, especially its medial lobe (mLH) which was previously shown to be food cue-responsive (*14*), whereas other related hormone receptors such as growth hormone receptor b (*ghrb*), were strongly expressed in the medial hypothalamus (MH) suggesting distinct functions **(Figure S4C-D).**

We further validated the co-expression of *tac1*, *ghra*, and their GABAergic identity *in situ* (**Figure 5A-B**). Quantitative analysis of *fosab* expression in this *tac1 and ghra*-positive cluster confirmed that it is significantly more active in the LH during voracious feeding as compared to food-deprivation (**Figure 5C**), which is consistent with the Act-seq data. Additionally, an increased number of *tac1*-expressing cells in the mLH were also quantified in this state (**Figure 5D**).

**Figure 5:**
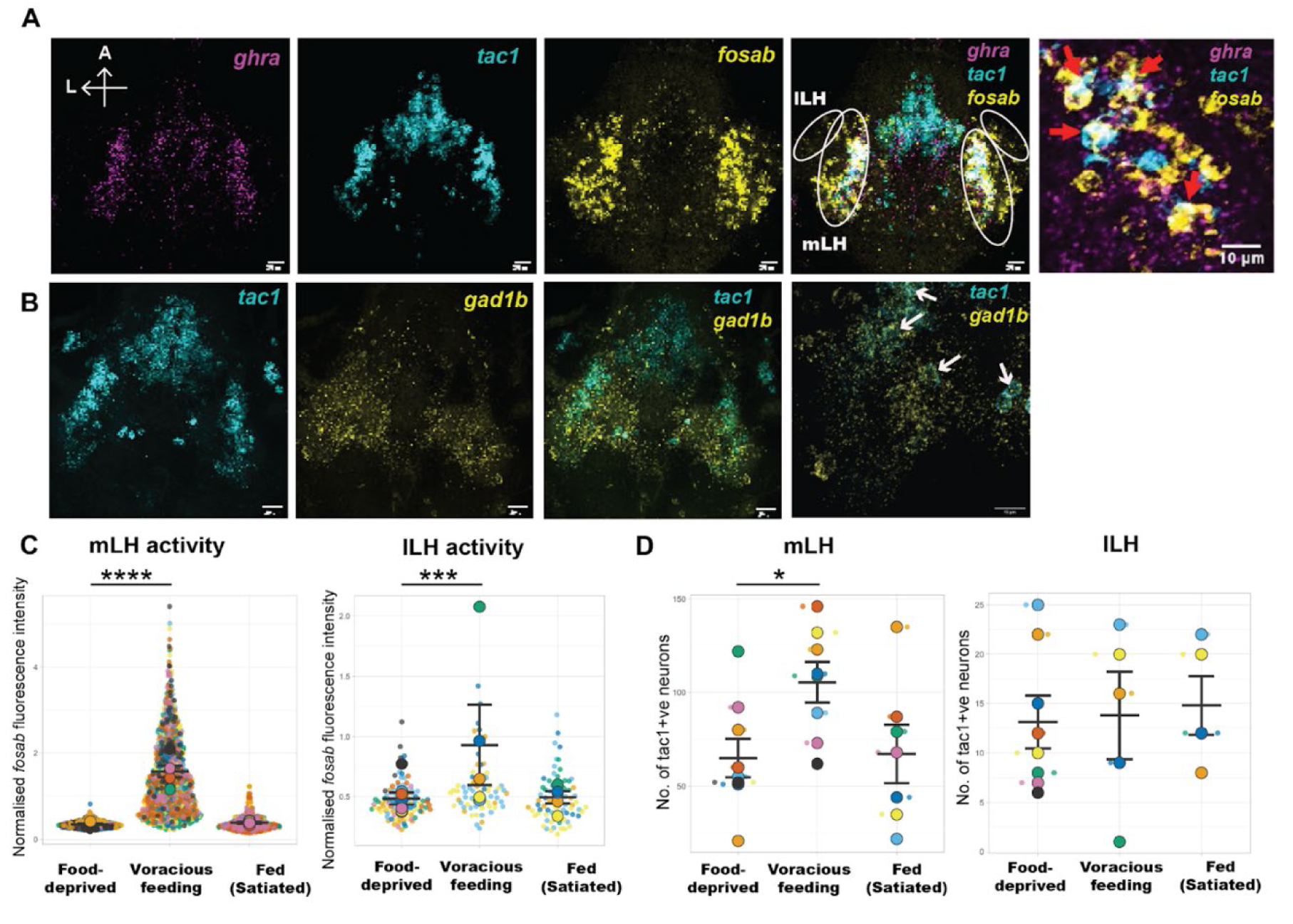
*Tac1/Ghr* neuronal cluster is dynamically regulated by feeding state. **A)** HCR validation of a GABAergic cluster (*tac1+ghra)* which is activated in voracious feeding conditions (Scale bar = 20 µm, magnified image scale bar = 10 µm). **B)** Validation of the GABAergic identity of *tac1* neurons (Scale bar = 20 µm). **C)** *tac1/ghra* neuronal activity is significantly higher in voracious feeding in medial lateral (p-value < 0.0001) and lateral lateral (p-value = 0.0002) hypothalamus. Statistical significance was calculated using Mann-Whitney test. Food-deprived: N (No. of fish) = 9, n (no. of *tac1* neurons): mLH = 585, lLH = 105; Voracious feeding: N = 8, n (mLH) = 844, n (lLH) = 69, Fed (Satiated): N = 7, n (mLH) = 470, n (lLH) = 74. **D)** Number of *tac1* neurons is significantly higher in voracious feeding in medial (p-value = 0.01) lateral hypothalamus. Statistical significance was calculated using Mann-Whitney test. Food-deprived: N (No. of fish) = 9, Voracious feeding: N = 8, Fed (Satiated): N = 7.

### Zebrafish lateral hypothalamic tachykinin neurons are sensitive to chemosensory food cues

To confirm that LH *tac1*-expressing neurons are responsive to food cues, we performed two-photon calcium imaging of the LH of 6 or 7 dpf larval zebrafish that express GCaMP6s in tac1 neurons (see Methods). *tac1* neurons can be found both in the MH and the LH. Upon delivery of E3 medium with food chemosensory cues (i.e., filtered E3 that had been incubated with AP-100 larval feed), as shown in **Figure 6A and B** calcium signals increased in *tac1* neurons (data from an example fish is shown in **Figure 6C-D**) in both medial and lateral parts of the hypothalamus (data from 19 fish is shown **Figure 6E-F**). Averaged calcium functions indicated that the overall response to food cues in both the medial and LH *tac1* neurons was elevated and sustained throughout the period of food cue delivery and persisted even after delivery of food cues ceased (**Figure 6G-I**). Hence, *tac1-*expressing neurons in both the MH and LH are activated by food chemosensory cues.

**Figure 6.**
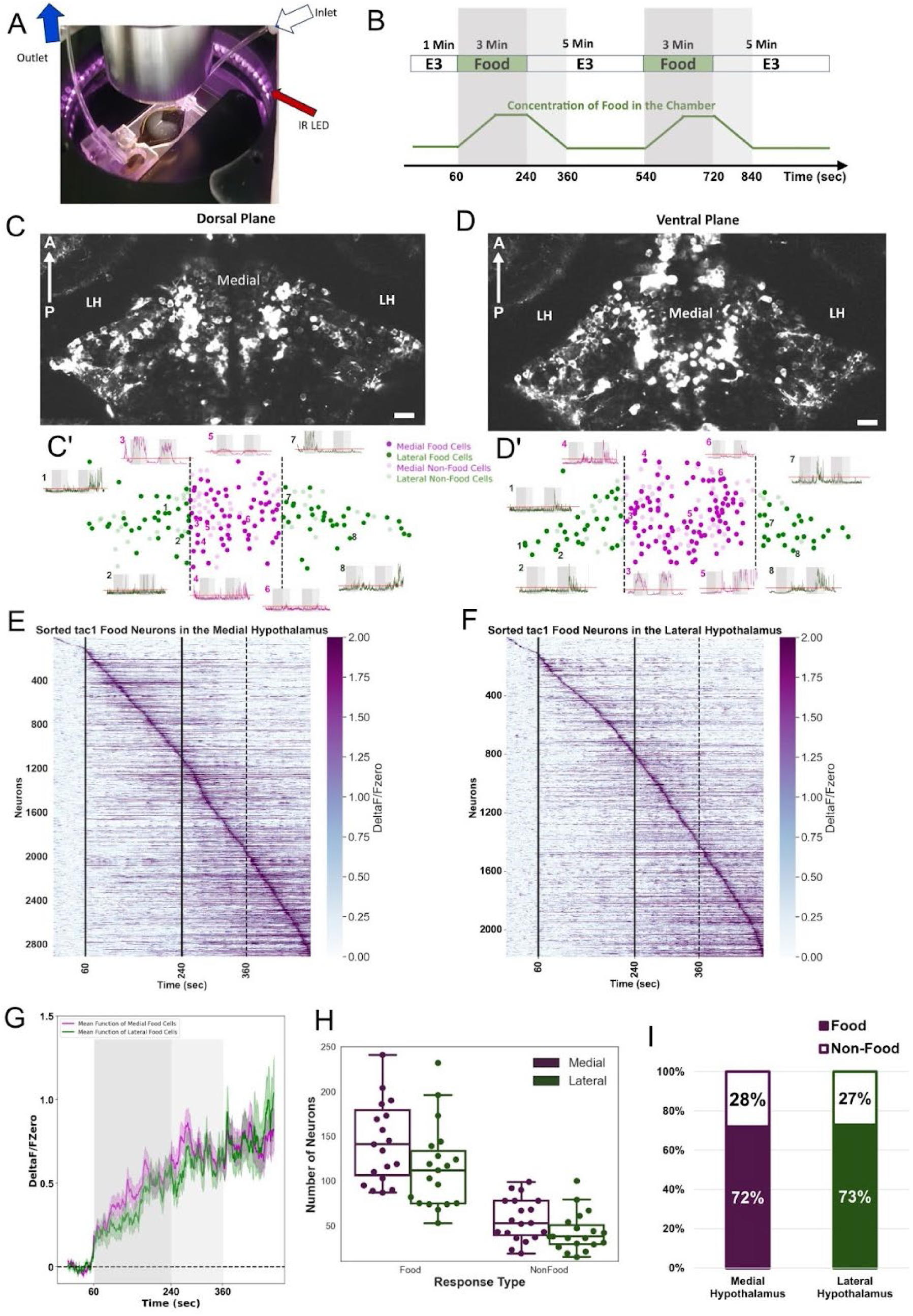
Tac1-positive hypothalamic neurons are responsive to food chemosensory cues. **A)** Photograph of the imaging setup showing the flow chamber in which the larva is mounted in agarose with the mouth region exposed and facing the inflow of food cues (E3 conditioned with AP100 larval feed) from the inlet (white arrow). **B)** Schematic of food cue delivery protocol. The fish is exposed to a continuous flow of E3 (white) until the stimulus is switched to food cues (dark gray) via a valve system. This protocol is repeated twice. The light gray area indicates the period where the food cues are likely still present but decreasing in concentration. The green function is the estimated food cues concentration as a function of time in the chamber estimated using a red dye as the infused medium following the same delivery program. **C**) The dorsal and **D)** ventral plane of the ventral hypothalamus of an example fish and spatial map (**C’& D’**) of food-cue responsive neurons (solid circles) versus non-responsive neurons (transparent circles). Calcium traces of representative food-responding neurons are displayed and numbered in **C’** and **D’**. Sorted heat maps show the average food cue-induced calcium activity of food-responding neurons in the medial **(E)** and lateral **(F)** hypothalamus across both stimulus periods. **(G)** Mean functions from the food-responsive neurons in the medial (magenta) versus the lateral hypothalamus (green) averaged across 19 fish. **H)** Number of food-responsive (left) vs non-responsive (right) neurons in the medial versus lateral hypothalamus per fish (n = 19 fish). **I)** Ratio of food versus non-food neurons in the medial versus the lateral hypothalamus. Scale bar = 20 μm.

### Modulation of zebrafish growth hormone and growth hormone receptor depends on feeding state

Previous work in zebrafish (and other fish species) has demonstrated that chronic (days to weeks) over-expression of GH can enhance food intake (*32*, *33*). The expression of *ghra* in “voracious feeding” neurons of the zebrafish LH indicates a potential role for GH in appetite regulation via the hypothalamic *tac1*/*ghra* circuit. Hence, we also characterized the dynamics of both GH (*gh1* in the pituitary) (**Figure 7A and B**) and its receptor (*ghra* in the LH) in zebrafish larvae (**Figure 7C**). Expression of the *gh1* transcript expression was the highest under voracious feeding and lowest in fed (satiated) larvae (**Figure 7B**). From *fosab* co-expression, we found that *gh1-*positive endocrine cells in the pituitary also had the highest activity in voraciously feeding larvae, suggestive of increased release of GH relative to that of food-deprived or fed (satiated) larvae (**Figure 7B**). Notably, *ghra* expression in the zebrafish LH was also regulated in the same fashion, increasing significantly during voracious feeding (**Figure 7C**). Together these results suggest that the transcription of genes encoding GH, and its receptor are dynamically modulated by hunger state and could potentially contribute to the acute regulation of food intake.

### Growth hormone modulates feeding state through potentiation of *tac1* neurons in zebrafish

To establish whether GH acutely regulates feeding via potentiation of *tac1*-expressing neurons, we treated 7 dpf larval zebrafish with human growth hormone (hGH). Interestingly, within 30 minutes of treatment, we observed an increase in food intake in fed (satiated) fish (**Figure 7D**). This increase in food intake was accompanied by an increase in *fosab* expression in *tac1* neurons (**Figure 7E**) suggesting that GH may directly act on *tac1* neurons in an acute manner through the GH receptors these cells express.

**Figure 7.**
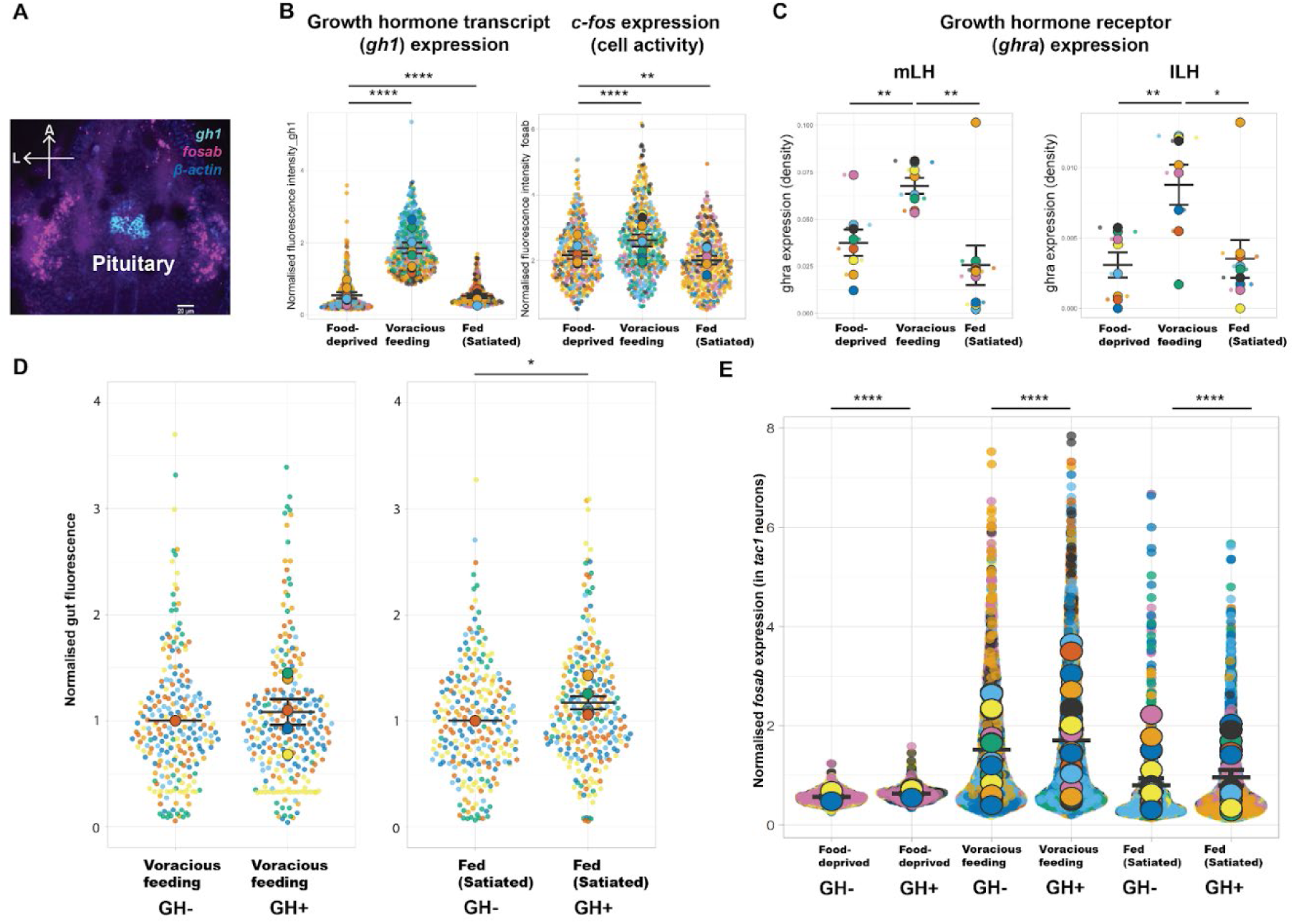
Growth hormone modulates feeding and neuronal activity in zebrafish. **A)** Growth hormone (*gh1)* expression in larval zebrafish pituitary (Scale bar = 20 µm) **B)** *gh1* transcript expression [Food-deprived: N (No. of fish) = 9, n (no. of endocrine cells) = 355; Voracious feeding: N = 8, n = 412, Fed (Satiated): N = 9, n = 343; p-value < 0.0001 for both comparisons] and activity of *gh1* expressing endocrine cells is modulated by feeding state with highest expression during voracious feeding (Food-deprived vs voracious feeding: p < 0.0001, Food-deprived vs fed: p = 0.001) **C)** Growth hormone receptor (*ghra)* expression in the medial lateral [Food deprived N = 8, Voracious feeding N = 8, Fed N = 9; mLH: Food-deprived vs voracious feeding: p-value = 0.003; Fed (Satiated) vs Voracious feeding: p-value = 0.005] and lateral lateral [Food deprived N = 8, Voracious feeding N = 8, Fed N = 9; lLH: Food-deprived vs voracious feeding: p-value = 0.004; Fed (Satiated) vs Voracious feeding: p-value = 0.03] is highest during voracious feeding . **D)** Human Growth Hormone (hGH) treatment results in increased feeding in satiated fish, as measured by gut fluorescence normalised to mean gut fluorescence of controls for each biological replicate [Voracious feeding (control): N (No. of fish) = 230, Voracious feeding (GH treated): N = 246, Fed (Satiated_control): N = 247, Fed (Satiated_GH treated): N = 261; p-value = 0.03, 6 replicates per condition] and **E)** increased activity in *tac1* neurons [Food-deprived_control: N (No. of fish) = 7, n (no. of tac1 neurons) = 1508; Food-deprived_GH treated: N = 6, n = 1471; Voracious feeding_ctrl: N = 13, n = 2336; Voracious feeding_GH treated: N = 19, n = 2528; Fed_ctrl (Satiated): N = 13, n = 1729; Fed_GH treated (Satiated): N = 13, n = 1941; p-value < 0.0001 for all three conditions). Statistical significance was calculated using Mann-Whitney test.

### *Tac1/Ghr* LH neurons are conserved in mammalian hypothalamus and dynamically modulated by feeding state

As described in **Figures 3** and **4**, a *Tac1/Ghr* expressing subpopulation was identified as a conserved GABAergic cluster. We validated the neuronal cell type in sections of the mouse brain with RNAScope, which demonstrated colocalization of *Ghr* transcripts (magenta) in *Tac1* neurons (green) (**Figure 8A**).

**Figure 8.**
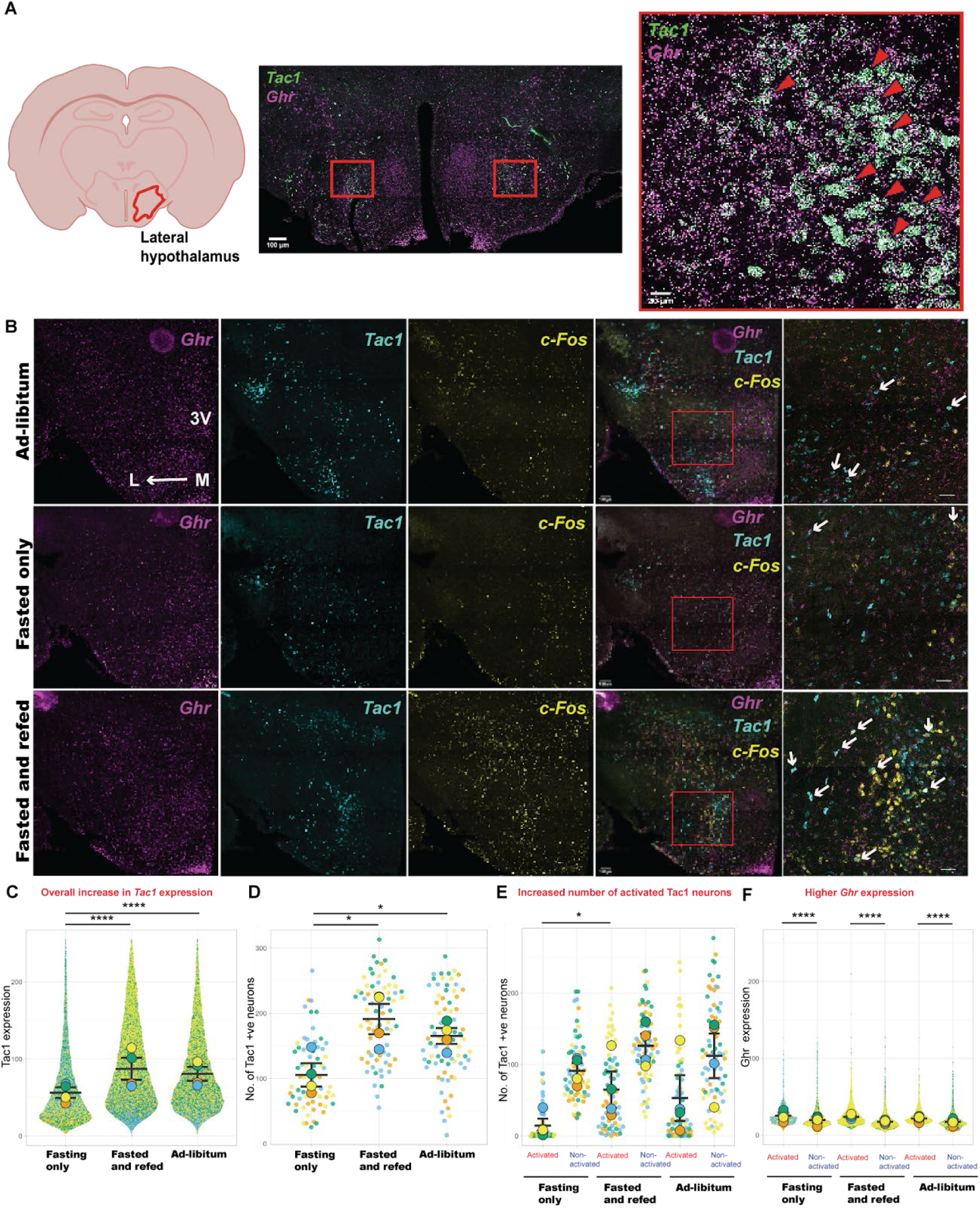
*Tac1/Ghr* neuronal cluster is conserved in mouse hypothalamus and is modulated by feeding state. **A)** RNAScope validation of *Tac1/Ghr* cluster in mouse brain sections shows co-expression of *Ghr* puncta (purple) in *Tac1* neurons (green) (Overview stitched image: Scale bar = 100 µm; Magnified image: Scale bar = 20 µm). **B)** Representative z-stack and stitched confocal images of mouse brain sections from “Fasted only”, “Fasted and refed” and “Ad-libitum” groups (Scale bar = 100 µm) with DAPI (blue), probes for growth hormone receptor (*Ghr)* (magenta), *Tac1* (cyan), *c-Fos* as an activity marker (yellow) and a merged image with an overlay of three channels **C)** Scatter plot showing a significant increase in expression of *Tac1* in Fasted and refed and Ad-libitum conditions (p <0.0001). D) Scatter plot showing a significant overall increase of *Tac1* neurons in “fasted and refed” state (p = 0.01) and “ad-libitum” state (p = 0.02). **E)** Scatter plot showing an increase in number of activated *Tac1* neurons in “fasted and refed” condition vs fasting only condition (p = 0.04). **F)** Scatter plot showing higher *Ghr* expression in activated *Tac1* neurons in all conditions (p < 0.0001). N = 4 brains (8-11 sections per brain) per condition. Statistical significance was calculated using Mann-Whitney test.

Next, we determined whether, similar to zebrafish, the mouse neurons are also modulated by feeding or nutrient status. Mice were subjected to a feeding paradigm in which five animals were assigned randomly to three groups that corresponded to the “food-deprived”, “voracious feeding” and “fed (satiated)” conditions in zebrafish: “fasted only” for animals deprived of food for 16 hours; “fasted and refed” when deprived of food for 16 hours followed by one hour of feeding; “ad-libitum” for animals provided unlimited food and water. Brain sections were collected that encompassed the entire lateral hypothalamus and RNAScope was used to detect *Tac1/Ghr* co-expressing neurons and *c-Fos* expression (**Figure 8B, Figure S5)** was used as a readout for neuronal activity. Notably, similar to zebrafish (**Figure 5**), we observed a significant increase in *Tac1* transcript expression (**Figure 8C**) and the number of LH *Tac1-*positive neurons (**Figure 8D**) in mice that were fasted and refed. To identify highly activated neurons, *c-Fos* expression levels were plotted as a histogram for each feeding condition with a stringent threshold of *c-Fos* > 30 corresponding to highly activated cells **(Figure S5)**. We observed a significant increase in the average number of activated *Tac1* neurons per brain in the “fast and refeeding” group relative to the “Fasting only” group (**Figure 8E**). Further, consistent with the zebrafish data, there was an overall increase in *Ghr* expression in activated *Tac1* neurons in the “Fasted and refed” (**Figure 8F**) condition suggesting that the *Tac1/Ghr* neuronal population is also dynamically affected by the feeding state in both expression and activity levels. Overall, our results suggest that the LH growth hormone-tachykinin circuit is dynamically responsive to feeding state and has a conserved role in appetite regulation both in zebrafish and mammals.

## DISCUSSION

The lateral hypothalamus has been established as a hunger center in vertebrate brains, while also regulating other diverse functions such as reward and motor activity (see detailed review by (*34*)). Different subpopulations of neurons even within the same region have distinct roles in appetite control (*20*). Hence, it is important to gain a comprehensive understanding of the molecular identities and functions of specific neuronal subtypes in response to feeding and hunger cues.

It was previously established in zebrafish that the LH and caudal hypothalamus (cH) are sensitive to internal hunger cues and food availability (*14*), but how these regions allow for nutrient sensing and control feeding was unclear. In this study, we used Act-seq to identify LH subpopulations that were more active in voracious feeding. By integrating mouse and zebrafish scRNAseq datasets, we demonstrate strong conservation especially of GABAergic subpopulations and identify a novel and conserved sub-population of LH neurons. This growth hormone-tachykinin cell type is dynamically regulated by the feeding state in both zebrafish and mice.

### Act-seq identifies novel food responsive sub-populations in zebrafish lateral hypothalamus and beyond

The spatial and molecular organisation of molecularly distinct cell types have been studied in the mammalian lateral hypothalamus by employing single-cell transcriptomics (*18*) or *in-situ* methods like EASI-FISH (*22*) on brain slices, albeit with a limited number of marker genes and coverage of the LH. Another study reported transcriptional changes in glutamatergic LH neurons during high-fat diet-induced obesity correlating with reduced appetite suppression, but did not pinpoint the specific subtypes involved (*21*).

By adopting the Act-seq method, we carried out a comprehensive single-cell profiling of whole zebrafish LH neuronal subpopulations that respond to an acute feeding stimulus, and identified neuronal clusters expressing well-known neuropeptides, receptors, and hormones which are dynamically regulated by feeding state. Zebrafish LH does not express some hallmarks of mammalian hypothalamic neurons such as Orexin (hypocretin) and Melanocortin-concentrating hormone (MCH) (*14*). However, several zebrafish neuronal subpopulations with GABAergic and glutamatergic identities co-express genes that were previously described in mice (*18*), and neuropeptides implicated in appetite regulation. Unlike glutamatergic clusters, all activated zebrafish GABAergic clusters had a corresponding mouse cluster, suggesting stronger conservation. Notably, the *tac1*/*ghra* GABAergic cluster, which is dynamically regulated by feeding state in both zebrafish and mice, was not previously reported in the mammalian dataset, but discovered from our comparative omics approaches.

We also identified other non-LH neurons that were activated during voracious feeding, such as in the cranial ganglia, cerebellar granule cells, and an *otpa*-expressing population. The cerebellum has been implicated in feeding regulation via multiple mechanisms including functional connectivity to regions such as the LH and by responding or influencing gut hormones and neurotransmitters (*35–37*). It is well-known that *otpa*-expressing cells are expressed in brain regions such as the ventral telencephalon, preoptic area, posterior tuberculum, hindbrain and spinal cord (*38*) which are all relevant regions for feeding regulation.

### Dynamic expression and activity of neuromodulatory LH populations

Of note, the expression of neuromodulators in the LH was dynamically regulated by the feeding state. We observed two broad classes of activity patterns. Neuronal subtypes positive for *tac1, penka, trh* and *vip*, or “voracious neurons”, showed the highest activity during voracious feeding and could be involved in upregulating feeding based on hunger cues. Other neuronal populations (e.g., expressing *ccka*, *penkb* and *pdyn*) were “food-responsive neurons” that were activated by food and remained active during satiety, suggesting that they signal the presence of food and subsequent satiety. The combined activity patterns of these broad cellular subtypes may be sufficient to explain the composite LH activity pattern reported in (*14*) in the absence or presence of food, while suggesting distinct functions for different neuropeptidergic populations. In mammals, anti-correlated activities between the ventromedial and lateral hypothalamus have been reported (*16*, *17*). In zebrafish, the lateral and caudal hypothalamus are similarly anti-correlated with a potential bidirectional crosstalk mechanism, although the detailed circuitry is still unknown (*14*). Of particular interest are the neuromodulator-receptor pairs expressed in the LH and cH. The identification of monoamine receptors in the zebrafish LH, including within activated clusters, suggests potential direct communication with the monoaminergic populations found in the cH, in the absence of any obvious neuronal projections connecting these regions.

### Integrative LH single-cell transcriptomics reveals cross-species conservation

Strong molecular parallels between other hypothalamic regions such as the mammalian paraventricular hypothalamus with the zebrafish neurosecretory preoptic area have previously been reported. However, the LH has remained less well-characterised due to its heterogeneous nature and the absence of key mammalian LH populations such as hypocretin and MCH expressing neurons. Based on developmental characterisation of *her5*-positive progenitors, Bloch et al (2019) (*19*) suggest that the zebrafish LH has a mesencephalic origin.

Here, we generated an integrated atlas of zebrafish and mouse LH neurons. By comparing integration efficacy and common marker genes, we identify stronger conservation of GABAergic clusters and highly conserved clusters such as the *th2/slc18a2* GABAergic population, along with the *tac1/ghra* population. We also identified cell types that are exclusive to zebrafish or mice. This dataset to our knowledge is the first comparative omics study of the zebrafish and mouse LH, paving the way for a better understanding of hypothalamic molecular and functional diversity over evolution.

One limitation of our analysis is that the LH is defined in our study by the *Tg (76A:Gal4FF)* transgenic line. Hence, it is possible that we may be missing out on certain zebrafish LH populations. However, given the broad expression of this transgenic line in the zebrafish LH and its detailed anatomical and functional characterisation in both larval and adult zebrafish (*14*, *15*, *39*) we are confident that our dataset provides ample coverage of this region. Secondly, the *Tg (76A:Gal4FF)* transgenic line labels a small subset of non-LH populations, which could be included in our analysis. We therefore validated key clusters using HCR. Unlike Bloch et al (2019) (*19*), we were unable to identify mesencephalic gene expression such as *her5* in this region, and instead found diencephalic marker genes such as *arxa, meis2, nkx2.2, otpa, shha, lhx5a, gbx2, barhl1b, fezf2 etc*.

### A growth hormone-tachykinin circuit proposed to regulate short term appetite

The expression of growth hormone receptor (*ghra*) in the GABAergic *tac1* population suggests that these neurons are responsive to growth hormone (GH). GH has been traditionally studied in the context of regulating peripheral metabolic processes. A small handful of studies have implicated GH in appetite regulation, however most of them only in the context of chronic exposure.

For example, chronic treatment with GH had an orexigenic effect in humans (*40*, *41*) and the same was observed in overexpression studies in mouse (*42*) and fish models (*32*, *33*, *43*). In children, chronic GH treatment increased appetite (*41*, *44*) while chronic overexpression of GH led to increased food intake in zebrafish (*43*) and coho salmon (*33*). In addition to feeding changes, chronic GH overexpression caused an increase in the expression of appetite regulating genes in the brain of coho salmon, suggesting that it acts directly on neuronal pathways regulating appetite. Currently, our understanding of central growth hormone pathways is limited to the well-characterized orexigenic agouti-related protein (AgRP) neurons (*45*) and not much is known about the involvement of other hypothalamic neurons expressing the growth hormone receptor. Also, since most studies involve chronic exposure to GH, it has been difficult to disentangle potential acute and direct effects of GH on appetite from longer-term metabolic effects.

Here, we show that mRNA levels of *gh1* and its receptor *ghra* are regulated by hunger state in the larval zebrafish brain. As expected, levels were highest in voracious feeding and lowest in the fed (satiated) state. These findings complement previous studies in humans and other fish models. For example, in human males, GH secretion was found to be significantly higher after short-term fasting (*46*) but decreased upon refeeding (*47*). Studies in common carp, goldfish, flounder, rainbow trout and salmon (*32*, *48*–*50*) reported that serum GH levels were highest in fasted and refed fish, with lowest circulating GH levels in fed (satiated) fish, though the functional consequences of such dynamic changes were not known.

Importantly, we found that a short-term (30-min) exposure to hGH was sufficient to both activate GABAergic *tac1/ghra* neurons and increase food intake in fed (satiated) fish. We thus propose that the GABAergic *tac1/ghra* neurons are a subpopulation through which GH induces its orexigenic effect.

Notably, we determined that an analogous GABAergic *Tac1/Ghr* cluster could be identified from a single-cell transcriptomics dataset of the mouse LH (*18*). We were able to validate the presence of this population in the mouse LH by *in-situ* hybridisation and found that, similar to zebrafish, the expression dynamics of growth hormone receptors on *Tac1* neurons as well as the activity of the *Tac1/Ghr* neurons were regulated by feeding state and enhanced during voracious feeding. The number of *Tac1*-expressing neurons and level of *Tac1* transcript expression were similarly enhanced during voracious feeding in both mice and zebrafish.

In mice, one hour of feeding was sufficient to significantly activate the *Tac1/Ghr* double positive population, suggesting that the growth hormone-tachykinin circuit is a conserved appetite regulating pathway even on a shorter time scale. Recently, GH has been reported to have an appetite-stimulating effect in mice on relatively short time scales (up to 24 hrs) (*45*). We speculate that at least part of these effects are mediated by the GABAergic *Tac1/Ghr* population.

To gain a better understanding of the newly identified growth hormone-tachykinin circuit, there is a need to identify upstream or downstream hormonal influences or synaptic connections and the corresponding signaling pathways. An interesting candidate to study would be ghrelin which is a well-established orexigenic peptide that is known to increase GH secretion via the growth hormone secretagogue receptor in mammals (*51*), though its role in zebrafish appetite control is still unclear (*52*, *53*). A recent study in mammals showed that expression of the growth hormone receptor in the brain is necessary for ghrelin’s acute actions on food intake (*54*). In another study, Ghr in the mouse LH was also shown to be necessary for food-seeking behaviour in food-restricted male mice (*55*). Since the *Tac1/Ghr* population is expressed widely in the LH, it would be interesting to examine whether these neurons participate in the ghrelin-GH axis of the brain to facilitate the orexigenic effects of ghrelin.

Overall, we have generated a comparative atlas of the zebrafish and mouse LH, demonstrating that the larval zebrafish model can serve as a discovery agent for dissecting genes, cell types and circuits that play conserved roles in integrative appetite control, including diverse dynamics across hunger and satiety states. Through our approach we have identified a novel and conserved hormonal circuit that may help regulate appetite and food intake in accordance with growth or metabolic needs.

## METHODS

### Zebrafish husbandry and maintenance

Embryos were kept in E3 medium in a 28°C incubator at 14:10 light/dark cycle. Larval and adult zebrafish were maintained at 14:10 light/dark cycle in the fish facility at Institute of Molecular and Cell Biology (IMCB). All transgenic lines and protocols used on zebrafish were approved by A*STAR Biomedical Research Council Institute of Animal Care and Use Committee (IACUC) #211612. All embryos were staged according to (*56*).

### Transgenic and mutant lines

Transgenic lines Tg(*76A:Gal4FF;UAS:GCaMP6s)*, *Tg(116A;Gal4FF:UAS:GFP*) and *Tg(ETvmat:GFP)* were previously described (*14*, *15*, *39*)*. Tg(tac1:Gal4FF)^c602^* is an unpublished line from the Halpern lab, generated by Jung-Hwa Choi using Gbait-hsp70:Gal4 donor DNA (*57*).Wild-type zebrafish from AB background were used for GH treatment, feeding and FISH experiments.

### Bulk transcriptomics

#### Sample preparation

Transgenic lines *Tg(76A:Gal4FF:UAS GCaMP6s)* and Tg(*116A:Gal4FF:UAS GFP)* were used to obtain lateral and caudal hypothalamic populations respectively. Whole brains were dissected from one month old transgenic zebrafish and eight brains were pooled for each of the four biological replicates. Dissected brains were transferred into Neurobasal medium (Thermofisher Scientific) with B27 supplement (Thermofisher Scientific). Dissociation was carried out with a Papain Dissociation System (Worthington Biochemical Corporation) using a modified protocol described by (*58*). Cell suspension was filtered using a 40 μm cell strainer to avoid cell clumps. DAPI (1:1000) was added to the cell samples prior to sorting in a BD FACSAria II SORP cell sorter and GCaMP6s/GFP+ve cells were collected in Neurobasal medium with B27 supplementation. Total RNA was extracted from sorted cell populations using Arcturus PicoPure RNA Isolation kit (Thermo Fisher Scientific) according to the manufacturer’s protocol.

#### Bulk RNA Sequencing

RNA samples were analyzed on Agilent Bioanalyzer for quality assessment with RNA Integrity Number (RIN) ranging from 7.5 to 9.5 with average of RIN 8.5. cDNA libraries were prepared using 0.5 ng of total RNA using the SMARTSeq v2 protocol (*59*) with the following modifications: 1. Addition of 20 µM TSO; 2. cDNA amplification of 16 cycles; 3. A total of 200 pg of cDNA was utilized with one-fifth of the standard reaction volume from the Illumina Nextera XT kit. Library fragment size distributions were assessed using the DNA High Sensitivity Reagent Kit on a Perkin Elmer Labchip system. Sequencing of all libraries was performed using an Illumina NovaSeq 6000 platform (Illumina, San Diego, CA, USA) with paired-end reads of 2 × 151 cycles (Illumina, San Diego, CA, USA) at a sequencing depth of ∼30 million reads per sample.

#### Alignment and filtering with raw reads

Quality control of the sequencing data was performed using FastQC and then the QC filtered FASTQ files were obtained. Paired-end reads were mapped to the Zebrafish reference genome (GRCz11) using STAR based on GRCz11 annotations. Feature counts per gene was obtained and used for follow-up statistical analysis. Differential expression analysis (Threshold: False Discovery Rate < 0.05 and log2(Fold change) > was performed using the DESeq2 package in R.

### RNA-Fluorescent in-situ hybridisation

#### Zebrafish feeding protocol

Starting from 5 dpf, larvae were fed excess paramecia once a day. At 7 dpf, these larval zebrafish were randomly divided into three groups: 1) “Food-deprived” where larvae were transferred into a new petri dish with fresh E3 medium and starved for four hours “Voracious feeding”, where larvae were transferred to a new petri dish with fresh E3, starved for four hours and fed excess paramecia for 30 minutes and 3) “Fed (satiated)”, where larvae were transferred to a new petri dish with fresh E3 media and fed excess paramecia for four hours. At the end of the experimental period, larvae of each group were fixed rapidly by funnelling them through a fine mesh sieve and dropping the sieve into an ice-cold tube of 4% Paraformaldehyde in PBS.

#### HCR^TM^ RNA-FISH and immunostaining

Hybridisation chain reaction fluorescent in-situ hybridisations (Molecular Instruments, HCR v3.0) were performed on dissected brains from 7 dpf zebrafish using a modified protocol described by (*60*, *61*). PFA fixed larvae were washed with PBS and dehydrated with serial dilutions of 25%, 50%, 75% and 100% methanol and stored in 100% methanol at -20°C until use. Larvae were then gradually rehydrated with serial dilutions of methanol and three PBS washes before being dissected. Dissected brains were dehydrated and stored in methanol at -20°C overnight. After methanol fixation, brains were rehydrated gradually with 75%, 50% and 25% methanol followed by five PBS washes. At the detection stage, larvae were pre-hybridised at 37°C with hybridisation buffer for 30 minutes followed by probe hybridisation at 37°C for 12-16 hours and excess probes were washed with 5X SSC washes. For the amplification stage, brains were incubated with an amplification buffer for 30 minutes at room temperature followed by amplification with hairpins overnight. The brains were washed with 5X SSC to remove excess hairpins and incubated with DAPI (1:1000) for 30 minutes followed by washes with 2X SSC thrice for five minutes each before imaging.

For immunostaining to validate the “*th2+slc18a2*” cluster, the protocol was similar up to rehydration of brains. Post rehydration, and five PBS washes, blocking was carried out with a 4% BSA solution in 2X SSC buffer for one hour at room temperature followed by incubation with primary antibodies, chicken anti-GFP (1:500, Abcam #ab13970) and anti-TH2 (1:500, Immunostar #22941) in 2X SSC overnight at 4°C. Prior to secondary antibody incubation, the brains were washed with 2X SSC buffer thrice for 10 minutes each. Alexa fluor 647 and 488 conjugated secondary antibodies (1:500, Invitrogen) were used and incubated for three hours at RT followed by three 10-minute 2X SSC washes. Brains were then incubated in DAPI as described above. The brains were mounted ventral to the coverslip and imaged on a LSM800 (Zeiss) microscope with the Zen software. Z-Stacks were taken at 2μm thickness using a 20x objective at a resolution of 1024 x 1024 pixels.

To quantify gene expression and activity of individual neurons, the multi-point tool was used to manually select neurons. Density of neuroreceptor expression was quantified using the Analyze particles function in Fiji/ImageJ and the following formula: Number of receptors/Area of ROI (LH/MH/cH). Fluorescence intensity was measured and normalized to β-actin signal. Significance testing was done using Graphpad prism (v9.4.1)

### Act-seq

Transgenic *Tg(76A:Gal4FF:UAS GCaMP6s)* zebrafish larvae were fed from 5 dpf onwards and at 7dpf, 50 larval zebrafish were randomly sorted into two groups: 1) “Food deprived” - starved for four hours 2) “Voraciously feeding” - starved for four hours and fed excess paramecia for 30 minutes. At the endpoint, larvae were rapidly euthanised in ice-cold water and brains were dissected within 40 minutes and transferred into ice-cold Neurobasal medium with B27 supplement. Dissociation was carried out with a Papain Dissociation System (Worthington Biochemical Corporation) and as described by (*58*) but with a shorter papain incubation time to avoid cell death. To avoid dissociation-induced activity and preserve hunger/feeding based neuronal activity patterns, actinomycin D was added at a concentration of 0.1ug/ml to the papain mixture to inhibit transcription (*25*). Larval brains were initially incubated in papain mixture for 10 minutes at 34°C, followed by gentle trituration with a p1000 pipette to ensure uniform dissociation and a final incubation for 5 minutes. FACS was performed as described earlier using a BD FACSAria II SORP cell sorter and GCaMP6s positive (LH) and negative (cells from all other brain regions) cells were collected in Neurobasal medium with B27 supplementation and 10% FBS as they were prone to cell death. Since the proportion of GCaMP6s positive LH cells is low as compared to non-LH cells in the brain, positive and negative populations were mixed at 50:50 ratio in both samples to ensure we captured a sufficient number of LH cells. Using the 10X Genomics Chromium Controller, about 10,000 cells were loaded for droplet encapsulation at a targeted cell recovery of about 6,000 cells. Single cell 3’ Gene Expression libraries were prepared using the 10X Genomics Chromium Next GEM Single Cell 3′ GEM, Library & Gel Bead Kit v3.1 and Chromium Dual Index Plate TT Set A Sample Index following the manufacturer’s instructions (10X Genomics, Pleasanton, CA, USA). The libraries were sequenced on an Illumina NovaSeq 6000 system (Illumina, San Diego, CA, USA) using indexed paired-end reads (2 × 151 bp), targeting a depth of approximately 50,000 reads per cell.

### Computational methods for single cell analysis

#### Alignment and filtering

The dataset was aligned to the zebrafish reference genome GRCz11 on PartekFlow using 10X Genomics CellRanger. Approximately 10,000 cells were sequenced, and quality filtering was performed using the following parameters: 200 < Number of expressed genes < 4000; maximum mitochondrial reads percent = 15. Quality filtering resulted in a total of 9031 cells (Food-deprived: 4412; Voraciously feeding: 4619). Following the standard Seurat pre-processing workflow, the data was normalized using the “LogNormalise” method which is a global-scaling normalization method, and a linear transformation was applied using the “ScaleData” function. Next, we performed a Principal component analysis (PCA) for linear dimensionality reduction. Elbow Plot was used to determine the dimensionality of the data and first 20 PCs were used for clustering using the “FindNeighbors” and “FindClusters” function at a clustering resolution of 0.5. Cells were divided into Gal4FF positive (average expression > 0.5) and negative (average expression <= 0.5) populations to identify lateral hypothalamic (LH) cells and differentiate them from the other populations.

#### Cluster annotation

LH cells were further divided into GABA and glutamatergic populations using the mean expression of *slc32a1* and *slc17a6a* respectively in each of the clusters. Both populations were then sub-clustered as described above at a cluster resolution of 0.5 giving rise to 11 GABAergic and 10 glutamatergic subclusters per population. Cluster markers were obtained by using the “FindAllMarkers” function with a log Fold change threshold of 0.25 and min.pct threshold of 0.25 using the Wilcoxon rank sum test. Amongst the top markers, clusters were annotated based on the co-expression of genes encoding for neuropeptides, receptors, transcription factors etc. either from the mammalian LHA dataset (*18*) or novel candidates of interest.

#### Cross species integration of mouse and zebrafish single-cell transcriptomic datasets

Before integration, BioMart was used to assign mouse orthologs to zebrafish genes and zebrafish gene names were converted to corresponding mouse gene names. In case of one-to-many orthologs, the zebrafish gene name was converted to mouse ortholog with highest % similarity. Integration was performed using Seurat canonical correlation analysis (CCA) on R (*31*) separately for the GABAergic and glutamatergic sub-populations of our zebrafish LH dataset and the previously published mouse LH dataset (*18*). We identified the features for integration using the “SelectIntegrationFeatures” function with n = 10,000. For the identification of integration anchors, “FindIntegrationAnchors” function was used with k.anchor = 50. For data integration, a k.weight of 40 was used to generate a batch corrected integrated dataset and this merged object was then used for downstream analyses. Integrated GABA and glutamatergic datasets were then sub-clustered at a resolution of 0.5 giving rise to 12 GABAergic and 16 glutamatergic clusters.

### Two-photon calcium imaging

Two-photon calcium imaging in larval zebrafish was conducted as described previously (*62*). In brief, the imaging was carried out using a Nikon A1RMP two-photon microscope with a 25x water dipping objective (NA=1.1) and a Toptica laser (FentoFibre ultra-920) with a fixed wavelength at 920 nm. The imaging speed was 1 frame-per-second on each scanned z plane (z = 2). Larval zebrafish were anesthetized in mivacurian (1.5mg/mL, Abcam, ab-143667) for 2 minutes and then embedded dorsal up in 2% low-melting agarose in E3 on a glass coverslip held in a Warner chamber (RC-26GLP, Warner Instruments). After the agarose was solidified, a wedge of the agarose with an angle of around 120 degrees was removed to expose the area surrounding the fish mouth. Odor stimulus (filtered Zeigler AP100) was delivered via a perfusion device (Warner VC-6M, Harvard Apparatus) controlled by TTL pulses from the Nikon DAQ board (National Instruments). The perfusion of food cues was delivered twice and each time lasted for 3 minutes and followed by 5 minutes of inter-trial-interval. Mouth movements of the fish were monitored using an InfraRed (IR) LED ring light and IR camera (Basler acA1440-220um USB3). All fish were verified to display mouth movements upon food cue exposure.

Nikon ND2 raw images were registered for motion correction using Suite2p built-in algorithms. The ROIs of each brain cell were automatically detected by Suite2p and further categorized by their response profiles using customized Python script for plotting the calcium response functions. The delta change in fluorescence intensity over the baseline (δF/F0) was calculated using the averaged fluorescence value from the 60 frames prior to the onset of the first odor stimulus as the F0. Subsequent fluorescence values were subtracted and divided by the F0 to obtain the ratio change. A neuron was considered to be responsive to food cues if its δF/F0 value exceeded 2 at any time during the food cues delivery time window as well as during the extended time window of 2 minutes after the perfusion device was turned off by the program (see the light gray area in **Fig. 6B**). Heatmaps and mean functions were obtained by averaging the δF/F0 ratio data from the 2 delivery time windows (see both of the dark and light gray areas).

### Growth hormone treatment and quantification of feeding in zebrafish

Larval zebrafish (7 dpf) were randomly assigned to six groups (25 fish per 10-cm petri dish); three feeding groups with a control and human growth hormone (hGH) treated set for each) “Food deprived (starved for 4 hours)”, “Voracious feeding (starved for 4 hours and fed with excess para for 30 minutes)” and “Fed (fed continuously for 4 hours with excess para)”. All the fish were fed in the morning for one hour and transferred into fresh E3 for the feeding assay as described earlier. Post feeding paradigm, the treatment groups were treated with 1 µM of hGH for 30 minutes while an equal volume of sterile water (which is the solvent used for GH) was added to the control groups. Post treatment, fish from “Voracious feeding” and “Fed (Satiated)” groups were fed for 30 minutes with excess paramecia labeled with 2.5 mg of lipid dye. At the end of this feeding period, fish were rapidly fixed in 4% paraformaldehyde and underwent overnight fixation at 4°C. Imaging of the fixed fish and quantification of gut fluorescence was performed as described in detail by (*63*). Gut fluorescence was used as a measure of paramecia consumed by each fish. Samples with fluorescence values exceeding 3 standard deviations from the mean of their respective groups were excluded from each experimental condition.

### Mouse experiments

#### Feeding paradigm

All experimental procedures on mice were approved by the institutional Animal Care and Use Committee (IACUC, protocol number: 211634) of the Agency for Science, Technology and Research (A*STAR) of Singapore. Mice were housed in a 12-hour light-dark cycle with access ad-libitum to water and standard chow diet (Altromin 1324 Maintenance diet). 8-week old male mice obtained from In Vivos Pte Ltd (Singapore) were randomly assigned to three groups (five animals per group): “Ad-libitum”: animals had access to unlimited food and water; “Fasting only”: animals were food deprived for 16 hours; “Fast and refed”: animals were food deprived for 16 hours and fed for one hour before being perfused.

#### RNAScope and analysis

Mice were deeply anesthetized at the endpoint of the feeding paradigm using ketamine and xylazine cocktail (ketamine: 150 mg/kg; xylazine: 10 mg/kg) and subsequently underwent transcardial perfusion with phosphate buffered saline (1X PBS) and then paraformaldehyde (4% PFA). Brains were extracted and subjected to overnight post fixation in 4% PFA at 4°C. Brains were then washed with 1X PBS and dehydration was carried out in 15% sucrose followed by 30% sucrose. Brains were embedded in Optimal cutting temperature compound (OCT) and cryosectioned coronally at 40 µm thickness. Sections were collected in 1X PBS and washed thrice with 1X PBS and mounted on Superfrost charged slides followed by a baking step to improve section adherence. Post-fixation was performed using pre-chilled 4% PFA followed by a dehydration process involving successive incubations in 50%, 70%, and 100% ethanol twice at room temperature. The sections were air-dried for 5 minutes and stored at -80°C until use. RNAscope™ Multiplex Fluorescent Reagent Kit v2 (ACDBio Inc.) was used for fluorescent in-situ hybridization (FISH). On the experiment day, slides were briefly thawed, and hydrogen peroxide was applied to each section for 10 minutes at room temperature, followed by two washes with 1X PBS. Target retrieval steps were then carried out by submerging the slides into boiled 1X detection target retrieval solution for 5 minutes and washed twice with 1X PBS. Sections were pre-treated with Protease Plus provided in the kit followed by probe hybridization for 2 hours at 40°C using the HybEZ^TM^ oven. Post-hybridization washes were done using a wash buffer to remove excess probes. Signal amplification steps included incubations with Amp1, Amp2, Amp3, followed by the application of peroxidase (HRP) conjugated C1, C2 or C3, depending on the channels used for the probes. Opal dyes (520, 620, and 690) were then applied to the slides to fluorescently label the targeted RNAs at 40°C. After final washes with a wash buffer, sections were stained with DAPI and fluoromount was applied before placing a coverslip and sealing it with nail polish. Sections were imaged using a LSM800 (Zeiss) microscope with the Zen software. Z-Stacks were taken at 2μm thickness using a 20x objective. For analysis, *Tac1* expressing neurons were selected and expression levels of *c-Fos* and *Ghr* were measured using custom written scripts on ImageJ. To identify highly active *Tac1* neurons in the LH, *c-Fos* expression levels were used to set a threshold. After identifying the mode of the data, the threshold was set at < 30 and cells expressing *c-Fos* levels above 30 were classified as highly active *Tac1* positive neurons.

## Acknowledgments

We acknowledge the funding support from the National Research Foundation Fellowship (NRF-NRFF13-2021-0003) awarded to Caroline Wee, and the Agency for Science, Technology and Research (A*STAR) Strategic Research Program (Brain-Body Initiative, iGrants call ID: #21718). We acknowledge core funding support from the Institute of Molecular and Cell Biology (IMCB), A*STAR, to Caroline Wee and Sarah Luo. We acknowledge core funding support from Singapore Immunology Network (SIgN), A*STAR and Singapore Food Story R&D Programme (W22W3D0003) grant support to Anand Kumar Andiappan. We acknowledge funding support from the National Medical Research Council Open Fund – Young Individual Research Grant (OF-YIRG) SC18/24-727015 awarded to Kimberle Shen. We would like to thank Nesha Afsha for her assistance in experiments. We would like to thank Shanshan Howland, Alicia Tay, Shihui Foo, Duan Kaibo and Nicholas Ang from the Singapore Immunology Network (SIgN) Immunomonitoring platform for their assistance with the transcriptomics studies and data curation and preliminary analysis. We would like to acknowledge the A*STAR Microscopy Platform (AMP) which was used for confocal microscopy. We would also like to thank Mohd Agus Bin Abdul Raman, Michael Bose and the IMCB Fish Facility staff for help in animal care.

## Funding

National Research Foundation Fellowship NRF-NRFF13-2021-0003 (CLW)

Agency for Science, Technology and Research (A*STAR) Strategic Research Program

Brain-Body Initiative, iGrants call ID: #21718 (CLW)

Singapore Food Story R&D programme: W22W3D0003 (AKA)

National Medical Research Council Open Fund-Young Individual Research Grant (OF-YIRG) SC18/24-727015 (KS)

## Author contributions

Conceptualization: VC, CLW

Methodology: VC, RKC, KS, NZ, CLW

Investigation: VC, RKC, NZ, GJD, SO, VYYT

Visualization: VC, RKC

Formal analysis: VC, RKC, AKA

Software: VC, RKC, ST Validation: GJD, ST

Resources: JHC, MEH, SXL, CLW

Supervision: AKA, WLC, SXL, CLW

Writing—original draft: VC, RKC, CLW

Writing—review & editing: VC, KS, NZ, JHC, MEH, AKA, SXL, CLW

## Data and materials availability

All data and code will be uploaded to https://github.com/CarolineWeeLab/LHtranscriptomics

## Supplementary Materials

**Fig. S1.**
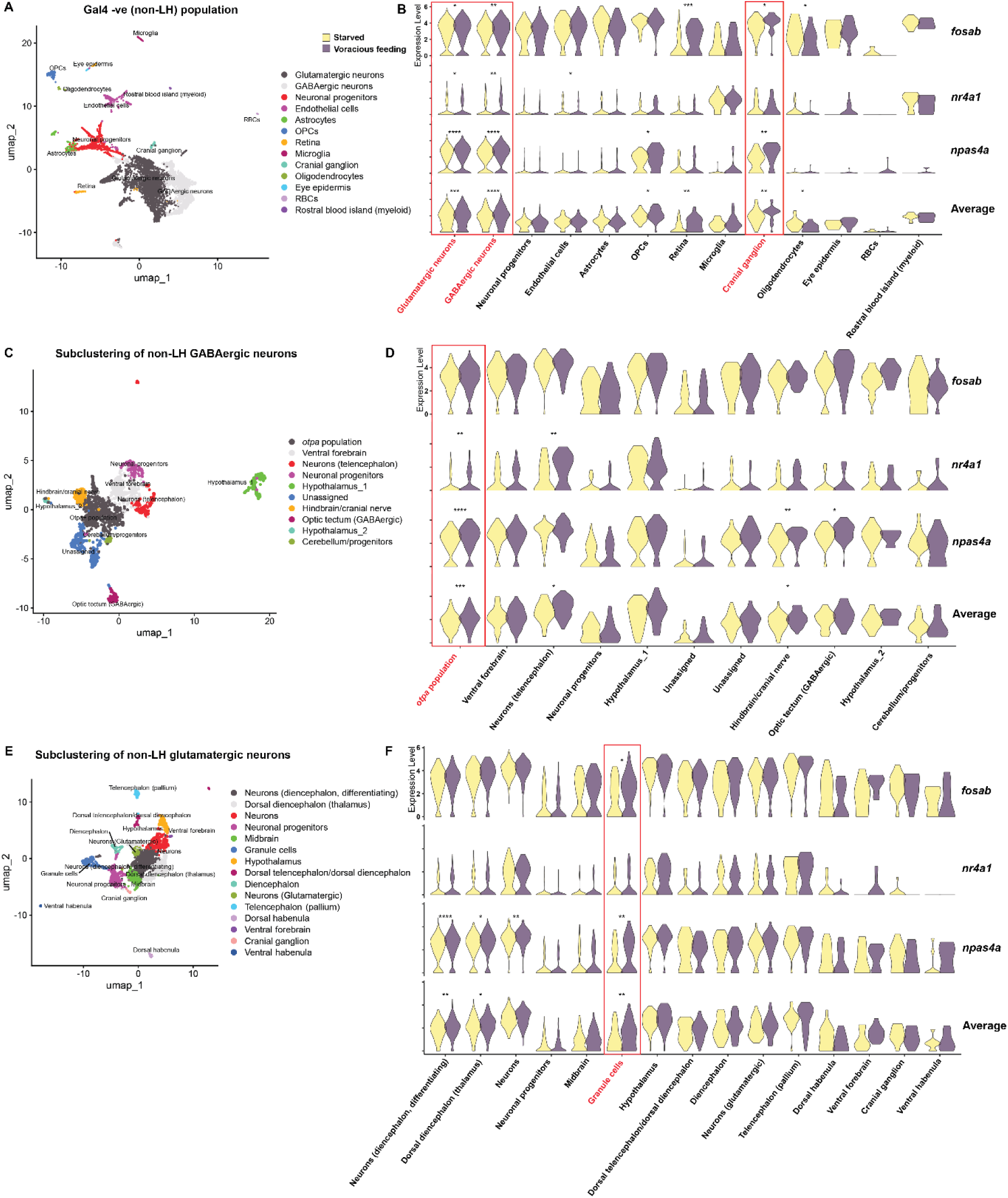
Act-seq analysis of non-Gal4-positive neurons. **A)** UMAP of all Gal4-negative (non LH) cells **C)** Sub-clustered non LH GABAergic neuronal population **E)** Sub-clustered non LH glutamatergic neuronal population. Violin plots showing expression levels of three IEGs, *fosab*, *nr4a1* and *npas4a* in **B)** all Gal4-negative (non-LH) cells **D)** non-LH GABAergic neuronal clusters **F)** non-LH glutamatergic neuronal clusters. Activated clusters (significant increase in expression of at least 2 IEGs) are highlighted in red.

**Fig. S2.**
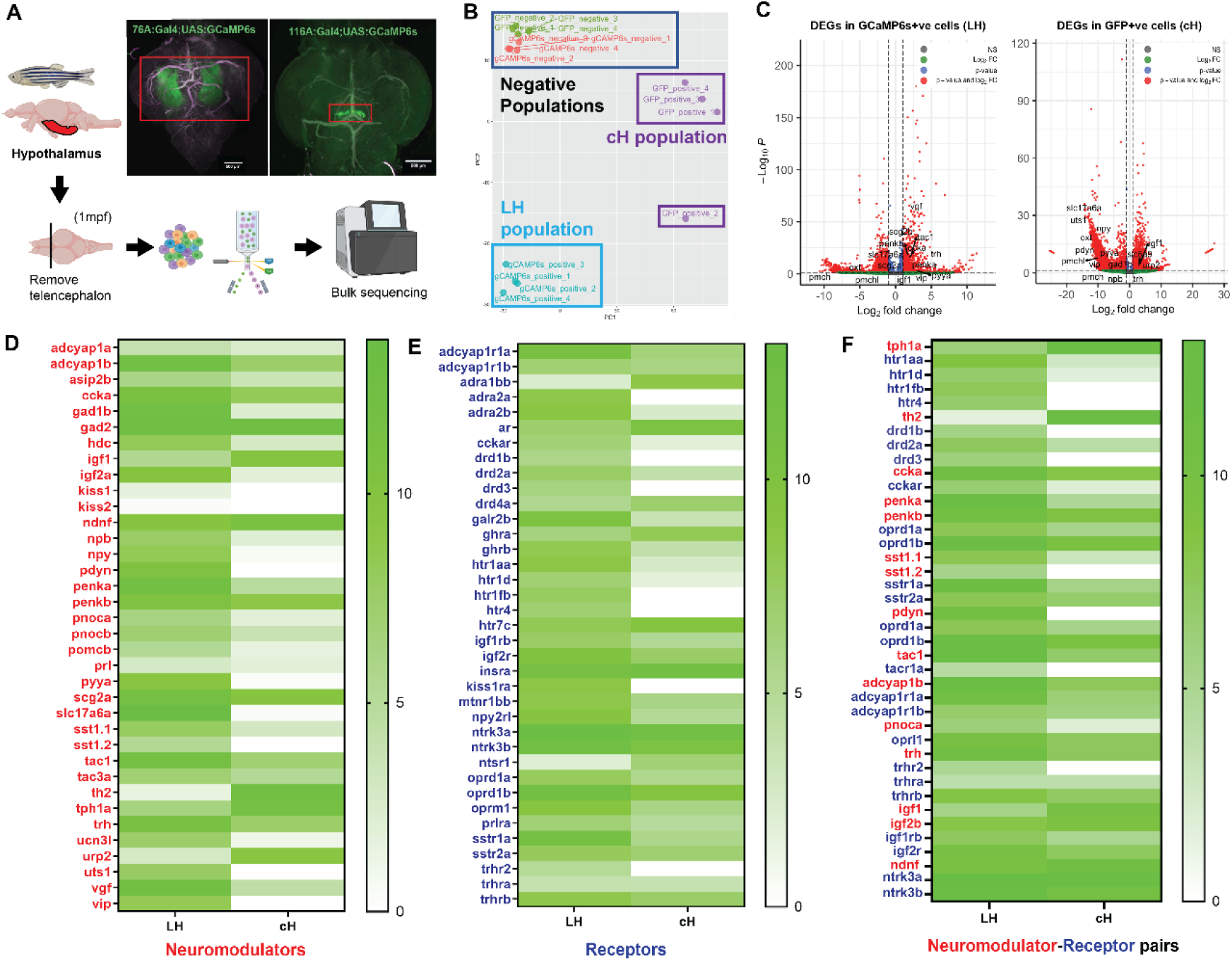
Bulk transcriptomics reveals differential expression of conserved neuropeptides and receptors in cH and LH. **A)** Schematic of sample collection and bulk transcriptomics. Scale bar = 500 µm **B)** Bulk transcriptomics PCA plot shows the clustering of LH and non-LH (GCaMP6s positive and negative respectively) and cH (GFP positive)and non-cH (GFP positive and negative respectively) **C)** Volcano plots showing differential expression of neuropeptides and receptors in LH (left) and cH (right) as compared to the rest of the brain. Heatmaps of normalized expression levels of **D)** conserved neuromodulators that are expressed differentially in cH and LH **E)** A subset of conserved neuroreceptors that are differentially expressed in both regions and **F)** neuromodulator (red) and receptor (blue) pairs identified in cH and LH as a potential mechanism for bi-directional communication.

**Fig. S3.**
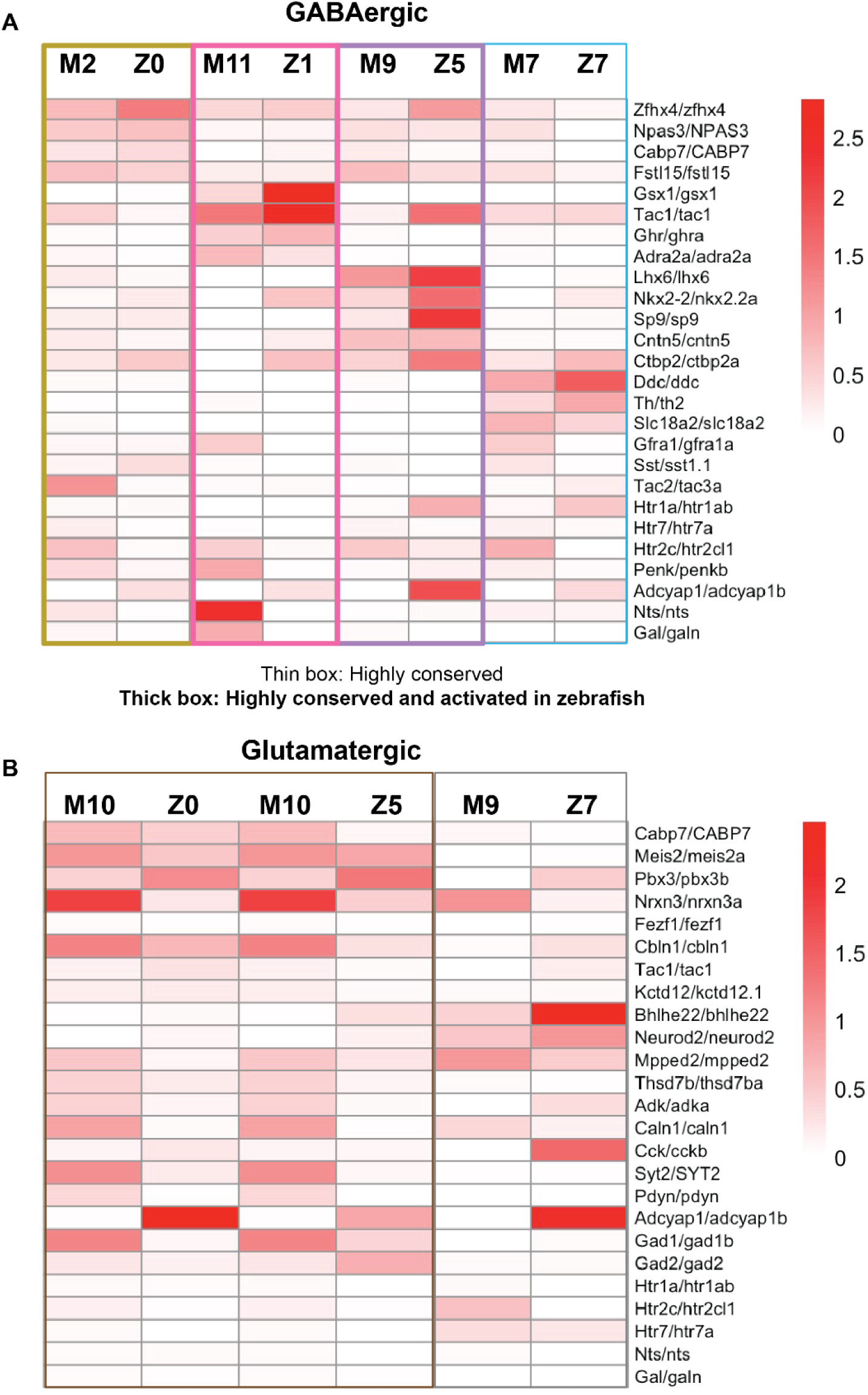
Inter species comparison of mouse and zebrafish LH. Heatmap visualization of common markers expressed in conserved **A)** GABAergic and **B)** glutamatergic clusters (note that M10^Glut^ is similar to both Z0^Glut^ and Z5^Glut^). Color code is the same as in **Figure 3**.

**Fig. S4.**
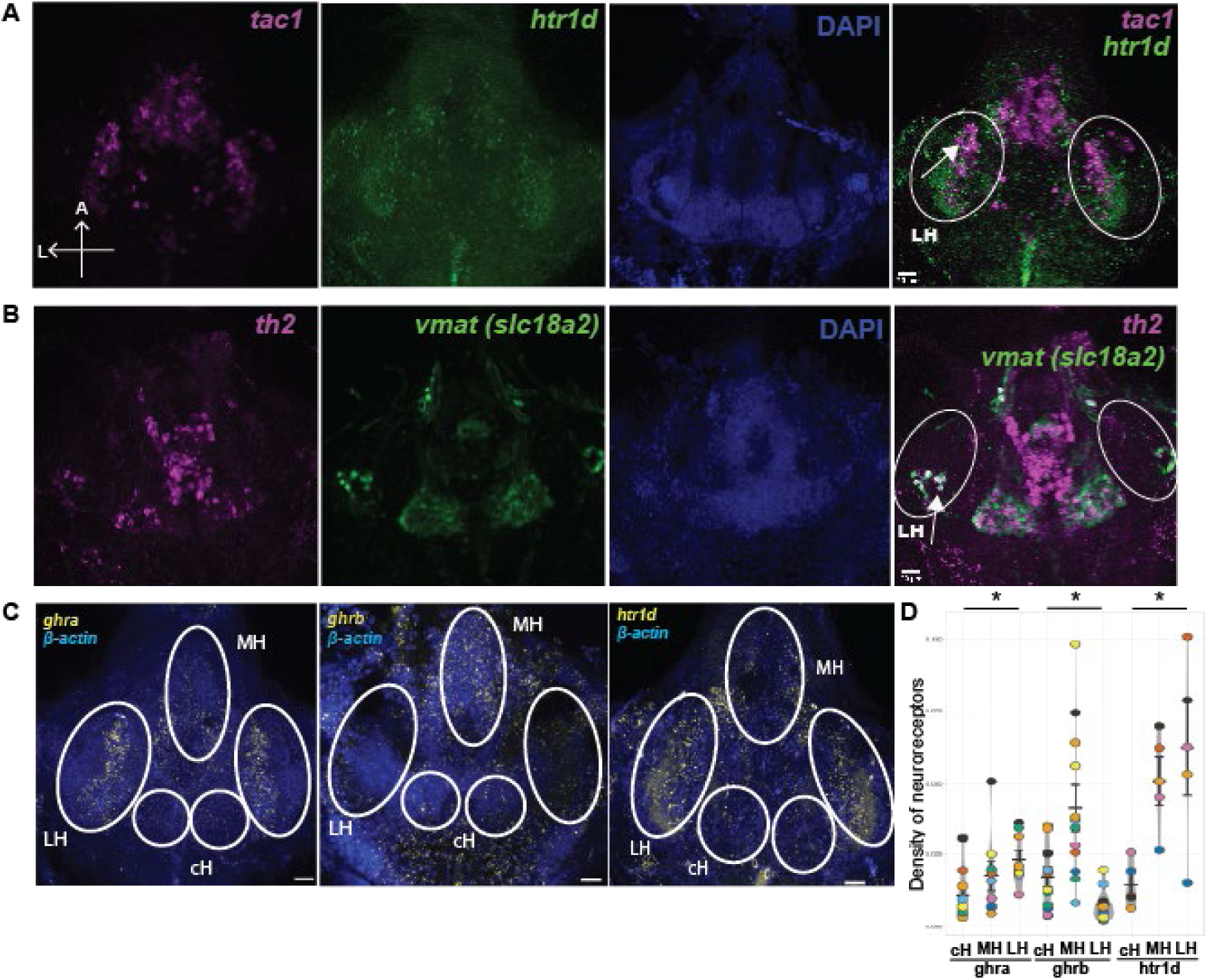
**Validation of additional clusters**. **A)** HCR validation of glutamatergic cluster, tac1+htr1d **B)** immunostaining to validate a conserved GABAergic cluster, expressing *th2* and *slc18a2* (*slc18a2* encodes for Vesicular monoamine transporter 2 or *Vmat2*). In ETvmat:GFP zebrafish brain, anti-Th2 antibody was used to label Th2-positive neurons and anti-GFP antibody was used to label vmat-positive neurons. **C)** Representative images showing expression of important neuroreceptors in 7dpf larval brains using HCR (*ghra, ghrb, htr1d*). Scale bar = 20 µm **D)** Scatter plot depicting density of neuroreceptors expressed in different hypothalamic regions (ghra: n =9 fish, p = 0.01; ghrb: n = 12, p = 0.01; htr1d: n = 5, p = 0.02). Significance testing was performed using unpaired t-tests. MH = Medial hypothalamus, LH = Lateral hypothalamus, cH = Caudal hypothalamus

**Fig. S5.**
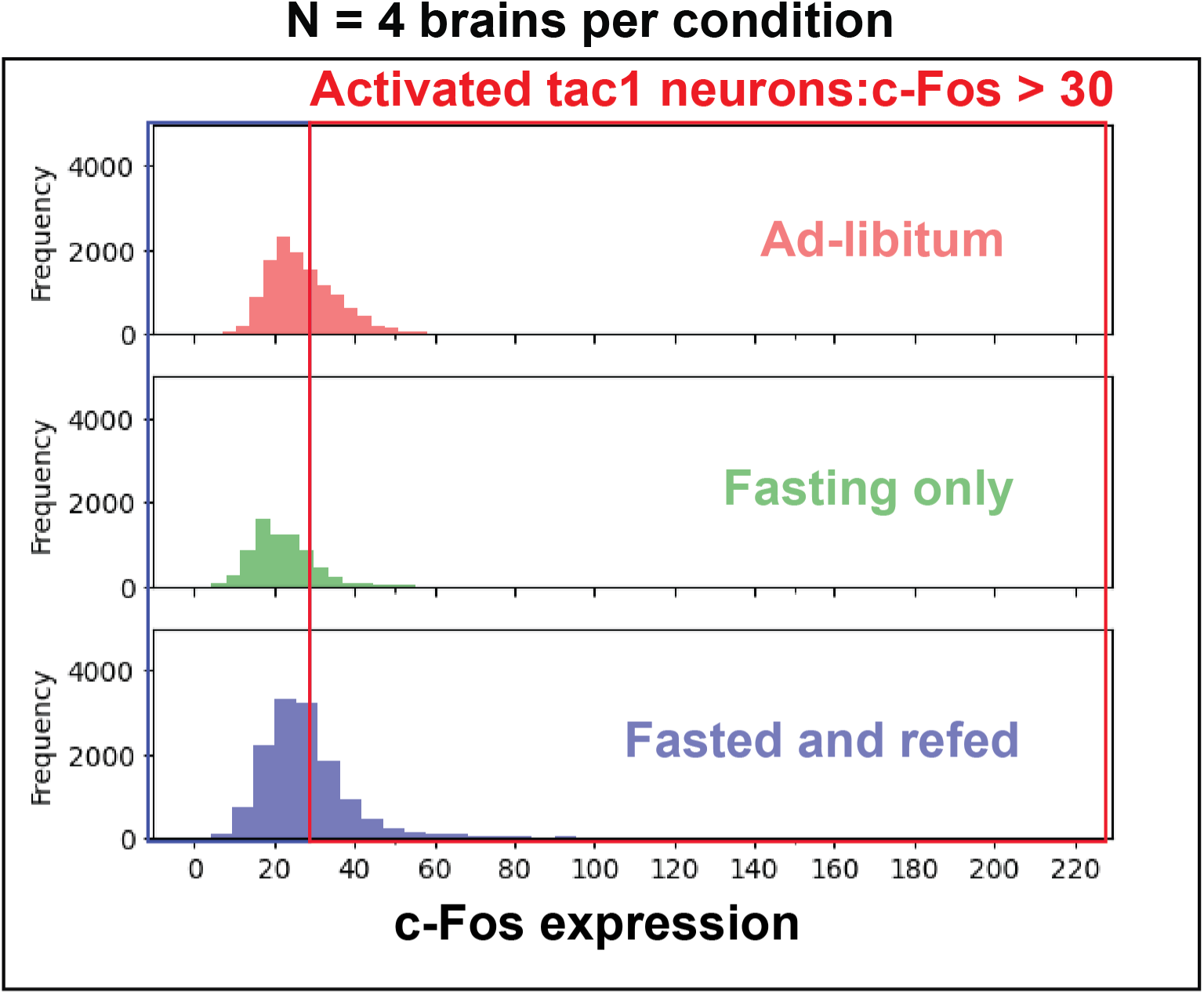
Histogram showing distribution of c-Fos in *Tac1* neurons in mouse brain sections in each feeding condition. 30 was chosen as a threshold to select for highly activated *Tac1* neurons.

---

**Data S1-S4 have been uploaded to** https://github.com/CarolineWeeLab/LHtranscriptomics

**Data S1**. Complete list of markers for zebrafish clusters (LH)

**Data S2**. Complete list of markers for zebrafish clusters (non-LH)

**Data S3**. Complete list of markers in mouse clusters (LH)

**Data S4**. Complete list of markers for integrated clusters (LH)

## References

1. L. M. Williams, Hypothalamic dysfunction in obesity. Proc. Nutr. Soc. 71, 521–533 (2012).

2. D. Kong, Q. Tong, C. Ye, S. Koda, P. M. Fuller, M. J. Krashes, L. Vong, R. S. Ray, D. P. Olson, B. B. Lowell, GABAergic RIP-Cre neurons in the arcuate nucleus selectively regulate energy expenditure. Cell 151, 645–657 (2012).

3. J. P. Thaler, C.-X. Yi, E. A. Schur, S. J. Guyenet, B. H. Hwang, M. O. Dietrich, X. Zhao, D. A. Sarruf, V. Izgur, K. R. Maravilla, H. T. Nguyen, J. D. Fischer, M. E. Matsen, B. E. Wisse, G. J. Morton, T. L. Horvath, D. G. Baskin, M. H. Tschöp, M. W. Schwartz, Obesity is associated with hypothalamic injury in rodents and humans. J. Clin. Invest. 122, 153–162 (2012).

4. C. Roger, A. Lasbleiz, M. Guye, A. Dutour, B. Gaborit, J.-P. Ranjeva, The Role of the Human Hypothalamus in Food Intake Networks: An MRI Perspective. Front Nutr 8, 760914 (2021).

5. A. W. Hetherington, S. W. Ranson, Hypothalamic lesions and adiposity in the rat. Anat. Rec. 78, 149–172 (1940).

6. B. K. Anand, J. R. Brobeck, Localization of a “Feeding Center” in the Hypothalamus of the Rat. Proc. Soc. Exp. Biol. Med. 77, 323–325 (1951).

7. P. M. Smith, A. V. Ferguson, Neurophysiology of hunger and satiety. Dev. Disabil. Res. Rev. 14, 96–104 (2008).

8. B. K. Anand, J. R. Brobeck, Hypothalamic control of food intake in rats and cats. Yale J. Biol. Med. 24, 123–140 (1951).

9. J. M. R. Delgado, B. K. Anand, Increase of food intake induced by electrical stimulation of the lateral hypothalamus. Am. J. Physiol. 172, 162–168 (1953).

10. J. R. Brobeck, S. Larsson, E. Reyes, A study of the electrical activity of the hypothalamic feeding mechanism. J. Physiol. 132, 358–364 (1956).

11. P. Teitelbaum, A. N. Epstein, The lateral hypothalamic syndrome: Recovery of feeding and drinking after lateral hypothalamic lesions. Psychol. Rev. 69, 74–90 (1962).

12. B. G. Hoebel, Hypothalamic Lesions by Electrocauterization: Disinhibition of Feeding and Self-Stimulation. Science 149, 452–453 (1965).

13. F. B. Krasne, General Disruption Resulting from Electrical Stimulus of Ventromedial Hypothalamus. Science 138, 822–823 (1962).

14. C. L. Wee, E. Y. Song, R. E. Johnson, D. Ailani, O. Randlett, J.-Y. Kim, M. Nikitchenko, A. Bahl, C.-T. Yang, M. B. Ahrens, K. Kawakami, F. Engert, S. Kunes, A bidirectional network for appetite control in larval zebrafish. Elife 8 (2019).

15. A. Muto, P. Lal, D. Ailani, G. Abe, M. Itoh, K. Kawakami, Activation of the hypothalamic feeding centre upon visual prey detection. Nat. Commun. 8, 15029 (2017).

16. Y. Oomura, K. Kimura, H. Ooyama, T. Maeno, M. Iki, M. Kuniyoshi, RECIPROCAL ACTIVITIES OF THE VENTROMEDIAL AND LATERAL HYPOTHALAMIC AREAS OF CATS. Science 143, 484–485 (1964).

17. Y. Oomura, H. Ooyama, T. Yamamoto, F. Naka, Reciprocal relationship of the lateral and ventromedial hypothalamus in the regulation of food intake. Physiol. Behav. 2, 97–115 (1967).

18. L. E. Mickelsen, M. Bolisetty, B. R. Chimileski, A. Fujita, E. J. Beltrami, J. T. Costanzo, J. R. Naparstek, P. Robson, A. C. Jackson, Single-cell transcriptomic analysis of the lateral hypothalamic area reveals molecularly distinct populations of inhibitory and excitatory neurons. Nat. Neurosci. 22, 642–656 (2019).

19. S. Bloch, M. Thomas, I. Colin, S. Galant, E. Machado, P. Affaticati, A. Jenett, K. Yamamoto, Mesencephalic origin of the inferior lobe in zebrafish. BMC Biol. 17, 22 (2019).

20. J. H. Jennings, G. Rizzi, A. M. Stamatakis, R. L. Ung, G. D. Stuber, The inhibitory circuit architecture of the lateral hypothalamus orchestrates feeding. Science 341, 1517–1521 (2013).

21. M. A. Rossi, M. L. Basiri, J. A. McHenry, O. Kosyk, J. M. Otis, H. E. van den Munkhof, J. Bryois, C. Hübel, G. Breen, W. Guo, C. M. Bulik, P. F. Sullivan, G. D. Stuber, Obesity remodels activity and transcriptional state of a lateral hypothalamic brake on feeding. Science 364, 1271–1274 (2019).

22. Y. Wang, M. Eddison, G. Fleishman, M. Weigert, S. Xu, T. Wang, K. Rokicki, C. Goina, F. E. Henry, A. L. Lemire, U. Schmidt, H. Yang, K. Svoboda, E. W. Myers, S. Saalfeld, W. Korff, S. M. Sternson, P. W. Tillberg, EASI-FISH for thick tissue defines lateral hypothalamus spatio-molecular organization. Cell 184, 6361–6377.e24 (2021).

23. L. Steuernagel, B. Y. H. Lam, P. Klemm, G. K. C. Dowsett, C. A. Bauder, J. A. Tadross, T. S. Hitschfeld, A. del Rio Martin, W. Chen, A. J. de Solis, H. Fenselau, P. Davidsen, I. Cimino, S. N. Kohnke, D. Rimmington, A. P. Coll, A. Beyer, G. S. H. Yeo, J. C. Brüning, HypoMap—a unified single-cell gene expression atlas of the murine hypothalamus. Nature Metabolism 4, 1402–1419 (2022).

24. R. Chen, X. Wu, L. Jiang, Y. Zhang, Single-Cell RNA-Seq Reveals Hypothalamic Cell Diversity. Cell Rep. 18, 3227–3241 (2017).

25. Y. E. Wu, L. Pan, Y. Zuo, X. Li, W. Hong, Detecting Activated Cell Populations Using Single-Cell RNA-Seq. Neuron 96, 313–329.e6 (2017).

26. R. E. Johnson, S. Linderman, T. Panier, C. L. Wee, E. Song, K. J. Herrera, A. Miller, F. Engert, Probabilistic Models of Larval Zebrafish Behavior Reveal Structure on Many Scales. Curr. Biol. 30, 70–82.e4 (2020).

27. J. Jordi, D. Guggiana-Nilo, E. Soucy, E. Y. Song, C. L. Wee, F. Engert, A high-throughput assay for quantifying appetite and digestive dynamics. Am. J. Physiol. Regul. Integr. Comp. Physiol., ajpregu.00225.2015 (2015).

28. J. H. Jennings, R. L. Ung, S. L. Resendez, A. M. Stamatakis, J. G. Taylor, J. Huang, K. Veleta, P. A. Kantak, M. Aita, K. Shilling-Scrivo, C. Ramakrishnan, K. Deisseroth, S. Otte, G. D. Stuber, Visualizing hypothalamic network dynamics for appetitive and consummatory behaviors. Cell 160, 516–527 (2015).

29. I. Shainer, E. Kuehn, E. Laurell, M. Al Kassar, N. Mokayes, S. Sherman, J. Larsch, M. Kunst, H. Baier, A single-cell resolution gene expression atlas of the larval zebrafish brain. Sci Adv 9, eade9909 (2023).

30. Z. V. Johnson, B. E. Hegarty, G. W. Gruenhagen, T. J. Lancaster, P. T. McGrath, J. T. Streelman, Cellular profiling of a recently-evolved social behavior in cichlid fishes. Nat. Commun. 14, 4891 (2023).

31. A. Butler, P. Hoffman, P. Smibert, E. Papalexi, R. Satija, Integrating single-cell transcriptomic data across different conditions, technologies, and species. Nat. Biotechnol. 36, 411–420 (2018).

32. C. Zhong, Y. Song, Y. Wang, T. Zhang, M. Duan, Y. Li, L. Liao, Z. Zhu, W. Hu, Increased food intake in growth hormone-transgenic common carp (Cyprinus carpio L.) may be mediated by upregulating Agouti-related protein (AgRP). Gen. Comp. Endocrinol. 192, 81–88 (2013).

33. J.-H. Kim, R. A. Leggatt, M. Chan, H. Volkoff, R. H. Devlin, Effects of chronic growth hormone overexpression on appetite-regulating brain gene expression in coho salmon. Mol. Cell. Endocrinol. 413, 178–188 (2015).

34. I. C. Alcantara, A. P. M. Tapia, Y. Aponte, M. J. Krashes, Acts of appetite: neural circuits governing the appetitive, consummatory, and terminating phases of feeding. Nat Metab 4, 836–847 (2022).

35. C. I. Iosif, Z. I. Bashir, R. Apps, J. Pickford, Cerebellar prediction and feeding behaviour. Cerebellum 22, 1002–1019 (2023).

36. M. Sader, G. D. Waiter, J. H. G. Williams, The cerebellum plays more than one role in the dysregulation of appetite: Review of structural evidence from typical and eating disorder populations, medRxiv (2022)p. 2022.04.14.22273867.

37. J. Zhao, M. Li, Y. Zhang, H. Song, K. M. von Deneen, Y. Shi, Y. Liu, D. He, Intrinsic brain subsystem associated with dietary restraint, disinhibition and hunger: an fMRI study. Brain Imaging Behav. 11, 264–277 (2017).

38. J. Eugenin von Bernhardi, D. Biechl, L. Miek, U. Herget, S. Ryu, M. F. Wullimann, A versatile transcription factor: Multiple roles of orthopedia a (otpa) beyond its restricted localization in dopaminergic systems of developing and adult zebrafish (Danio rerio) brains. J. Comp. Neurol. 530, 2537–2561 (2022).

39. L. Wen, W. Wei, W. Gu, P. Huang, X. Ren, Z. Zhang, Z. Zhu, S. Lin, B. Zhang, Visualization of monoaminergic neurons and neurotoxicity of MPTP in live transgenic zebrafish. Dev. Biol. 314, 84–92 (2008).

40. Y. E. Snel, R. J. Brummer, M. E. Doerga, P. M. Zelissen, H. P. Koppeschaar, Energy and macronutrient intake in growth hormone-deficient adults: the effect of growth hormone replacement. Eur. J. Clin. Nutr. 49, 492–500 (1995).

41. J. Blissett, G. Harris, J. Kirk, Effect of growth hormone therapy on feeding problems and food intake in children with growth disorders. Acta Paediatr. 89, 644–649 (2000).

42. M. Bohlooly-Y, B. Olsson, C. E. G. Bruder, D. Lindén, K. Sjögren, M. Bjursell, E. Egecioglu, L. Svensson, P. Brodin, J. C. Waterton, O. G. P. Isaksson, F. Sundler, B. Ahrén, C. Ohlsson, J. Oscarsson, J. Törnell, Growth hormone overexpression in the central nervous system results in hyperphagia-induced obesity associated with insulin resistance and dyslipidemia. Diabetes 54, 51–62 (2005).

43. M. G. Meirelles, B. F. Nornberg, T. L. R. da Silveira, M. T. Kütter, C. G. Castro, J. R. B. Ramirez, V. Pedrosa, L. A. Romano, L. F. Marins, Growth Hormone Overexpression Induces Hyperphagia and Intestinal Morphophysiological Adaptations to Improve Nutrient Uptake in Zebrafish. Front. Physiol. 12, 723853 (2021).

44. M. Yackobovitch-Gavan, G. Gat-Yablonski, B. Shtaif, S. Hadani, S. Abargil, M. Phillip, L. Lazar, Growth hormone therapy in children with idiopathic short stature - the effect on appetite and appetite-regulating hormones: a pilot study. Endocr. Res. 44, 16–26 (2019).

45. I. C. Furigo, P. D. S. Teixeira, G. O. de Souza, G. C. L. Couto, G. G. Romero, M. Perelló, R. Frazão, L. L. Elias, M. Metzger, E. O. List, J. J. Kopchick, J. Donato, Growth hormone regulates neuroendocrine responses to weight loss via AgRP neurons. Nat. Commun. 10, 1–11 (2019).

46. M. L. Hartman, J. D. Veldhuis, M. L. Johnson, M. M. Lee, K. G. Alberti, E. Samojlik, M. O. Thorner, Augmented growth hormone (GH) secretory burst frequency and amplitude mediate enhanced GH secretion during a two-day fast in normal men. J. Clin. Endocrinol. Metab. 74, 757–765 (1992).

47. M. L. Hartman, Fasting-induced enhancement of pulsatile growth hormone (GH) secretion is rapidly abolished by refeeding. Prog 72nd Meet Endocrine Soc (1990).

48. L. F. Canosa, S. Unniappan, R. E. Peter, Periprandial changes in growth hormone release in goldfish: role of somatostatin, ghrelin, and gastrin-releasing peptide. Am. J. Physiol. Regul. Integr. Comp. Physiol. 289, R125–33 (2005).

49. E. N. Fuentes, I. E. Einarsdottir, J. A. Valdes, M. Alvarez, A. Molina, B. T. Björnsson, Inherent growth hormone resistance in the skeletal muscle of the fine flounder is modulated by nutritional status and is characterized by high contents of truncated GHR, impairment in the JAK2/STAT5 signaling pathway, and low IGF-I expression. Endocrinology 153, 283–294 (2012).

50. J.-C. Gabillard, K. Yao, M. Vandeputte, J. Gutierrez, P.-Y. Le Bail, Differential expression of two GH receptor mRNAs following temperature change in rainbow trout (Oncorhynchus mykiss). J. Endocrinol. 190, 29–37 (2006).

51. T. D. Müller, R. Nogueiras, M. L. Andermann, Z. B. Andrews, S. D. Anker, J. Argente, R. L. Batterham, S. C. Benoit, C. Y. Bowers, F. Broglio, F. F. Casanueva, D. D’Alessio, I. Depoortere, A. Geliebter, E. Ghigo, P. A. Cole, M. Cowley, D. E. Cummings, A. Dagher, S. Diano, S. L. Dickson, C. Diéguez, R. Granata, H. J. Grill, K. Grove, K. M. Habegger, K. Heppner, M. L. Heiman, L. Holsen, B. Holst, A. Inui, J. O. Jansson, H. Kirchner, M. Korbonits, B. Laferrère, C. W. LeRoux, M. Lopez, S. Morin, M. Nakazato, R. Nass, D. Perez-Tilve, P. T. Pfluger, T. W. Schwartz, R. J. Seeley, M. Sleeman, Y. Sun, L. Sussel, J. Tong, M. O. Thorner, A. J. van der Lely, L. H. T. van der Ploeg, J. M. Zigman, M. Kojima, K. Kangawa, R. G. Smith, T. Horvath, M. H. Tschöp, Ghrelin. Mol Metab 4, 437–460 (2015).

52. K. Guan, C. Shan, A. Guo, X. Gao, X. Li, Ghrelin regulates hyperactivity-like behaviors via growth hormone signaling pathway in zebrafish (Danio rerio). Front. Endocrinol. (Lausanne) 14, 1163263 (2023).

53. X. Li, J. He, W. Hu, Z. Yin, The essential role of endogenous ghrelin in growth hormone expression during zebrafish adenohypophysis development. Endocrinology 150, 2767–2774 (2009).

54. F. Wasinski, F. Barrile, J. A. B. Pedroso, P. G. F. Quaresma, W. O. Dos Santos, E. O. List, J. J. Kopchick, M. Perelló, J. Donato, Ghrelin-induced Food Intake, but not GH Secretion, Requires the Expression of the GH Receptor in the Brain of Male Mice. Endocrinology 162 (2021).

55. M. R. Tavares, W. O. Dos Santos, I. C. Furigo, E. O. List, J. J. Kopchick, J. Donato Jr, Growth hormone receptor in lateral hypothalamic neurons is required for increased food-seeking behavior during food restriction in male mice. J. Neurosci. 44 (2024).

56. C. B. Kimmel, W. W. Ballard, S. R. Kimmel, B. Ullmann, T. F. Schilling, Stages of embryonic development of the zebrafish. Dev. Dyn. 203, 253–310 (1995).

57. Y. Kimura, Y. Hisano, A. Kawahara, S.-I. Higashijima, Efficient generation of knock-in transgenic zebrafish carrying reporter/driver genes by CRISPR/Cas9-mediated genome engineering. Sci. Rep. 4, 1–7 (2014).

58. B. Raj, J. A. Farrell, J. Liu, J. El Kholtei, A. N. Carte, J. Navajas Acedo, L. Y. Du, A. McKenna, Đ. Relić, J. M. Leslie, A. F. Schier, Emergence of Neuronal Diversity during Vertebrate Brain Development. Neuron 108, 1058–1074.e6 (2020).

59. S. Picelli, O. R. Faridani, A. K. Björklund, G. Winberg, S. Sagasser, R. Sandberg, Full-length RNA-seq from single cells using Smart-seq2. Nat. Protoc. 9, 171–181 (2014).

60. H. M. T. Choi, C. R. Calvert, N. Husain, D. Huss, J. C. Barsi, B. E. Deverman, R. C. Hunter, M. Kato, S. M. Lee, A. C. T. Abelin, A. Z. Rosenthal, O. S. Akbari, Y. Li, B. A. Hay, P. W. Sternberg, P. H. Patterson, E. H. Davidson, S. K. Mazmanian, D. A. Prober, M. van de Rijn, J. R. Leadbetter, D. K. Newman, C. Readhead, M. E. Bronner, B. Wold, R. Lansford, T. Sauka-Spengler, S. E. Fraser, N. A. Pierce, Mapping a multiplexed zoo of mRNA expression. Development 143, 3632–3637 (2016).

61. H. M. T. Choi, M. Schwarzkopf, M. E. Fornace, A. Acharya, G. Artavanis, J. Stegmaier, A. Cunha, N. A. Pierce, Third-generation in situ hybridization chain reaction: multiplexed, quantitative, sensitive, versatile, robust. Development 145 (2018).

62. J. S. M. Chia, E. S. Wall, C. L. Wee, T. A. J. Rowland, R.-K. Cheng, K. Cheow, K. Guillemin, S. Jesuthasan, Bacteria evoke alarm behaviour in zebrafish. Nat. Commun. 10, 3831 (2019).

63. R.-K. Cheng, J. X. M. Tan, K. X. Chua, C. J. X. Tan, C. L. Wee, Osmotic Stress Uncovers Correlations and Dissociations Between Larval Zebrafish Anxiety Endophenotypes. Front. Mol. Neurosci. 15, 900223 (2022).

